# Cold-triggered induction of ROS- and raffinose-related metabolism in freezing-sensitive taproot tissue of sugar beet

**DOI:** 10.1101/2021.04.12.439442

**Authors:** Isabel Keller, Christina Müdsam, Cristina Martins Rodrigues, Dominik Kischka, Wolfgang Zierer, Uwe Sonnewald, Karsten Harms, Olaf Czarnecki, Karin Fiedler-Wiechers, Wolfgang Koch, H. Ekkehard Neuhaus, Frank Ludewig, Benjamin Pommerrenig

**Author notes:** Corresponding Author: Dr. Benjamin Pommerrenig, Plant Physiology, University of Kaiserslautern, Paul-Ehrlich-Str 22, 67663 Kaiserslautern, Germany.

## Abstract

Sugar beet (*Beta vulgaris subsp. vulgaris*) is the exclusive source of sugar in the form of sucrose in temperate climate zones. There, sugar beet is grown as an annual crop from spring to autumn because of the damaging effect of freezing temperatures to taproot tissue. Natural and breeded varieties display variance in the degree of tolerance to freezing temperatures and genotypes with elevated tolerance to freezing have been isolated. Here we compare initial responses to frost between genotypes with either low and high winter survival rates. The selected genotypes differed in the severity of frost injury. We combined transcriptomic and metabolite analyses of leaf- and taproot tissues from such genotypes to elucidate mechanisms of the early freezing response and to dissect genotype- and tissue-dependent responses. Freezing temperatures induced drastic downregulation of photosynthesis-related genes in leaves but upregulation of genes related to minor carbohydrate metabolism, particularly of genes involved in raffinose metabolism in both, leaf and taproot tissue. In agreement with this, it has been revealed that raffinose and the corresponding intermediates, inositol and galactinol, increased markedly in these tissues. We found that genotypes with improved tolerance to freezing, showed higher accumulation of raffinose in a defined interior region within the upper part of the taproot, the pith, representing the tissue most susceptible to freeze damages. This accumulation was accompanied by specific upregulation of raffinose synthesizing enzymes in taproots, suggesting a protective role for raffinose and its precursors for freezing damage in sugar beet.

## INTRODUCTION

In temperate climate zones (Europe and North America), sugar beet (*Beta vulgaris subsp. vulgaris*) is the exclusive source of sugar (sucrose) for the food industry and a source for bio-energy generation. Sugar beet taproots are able to accumulate sucrose to up to nearly 20% of their fresh weight at maturity (Dohm et al., 2013) and provide about 30% of the total sugar produced worldwide (Y.-F. Zhang et al., 2017). Owing to its biennial lifestyle, the plant forms the huge sucrose-storing taproot during the first year of its life cycle. The stored sugar is used to fuel the outgrowth of a flowering seed stalk in the reproductive phase in the second year of growth (Chen et al., 2016). Induction of flowering, however, requires a prolonged exposure to cold (between 5-20 weeks at 4-15°C), known as vernalization (Abo-Elwafa et al., 2006; Kockelmann et al., 2010). During vernalization, shoots and taproots prepare for metabolic and functional rearrangements resulting in a switch of their source and sink identities, where shoot metabolism depends on carbon supply from the taproot to allow outgrowth of the seed stalk (Rodrigues et al., 2020). Successful vernalization of sugar beet plants can only occur at low, but above-zero temperatures. This is because, despite the high accumulation of sugars, which are known to stabilize membranes and protect against freezing-induced damages (Anchordoguy et al., 1987; Pommerrenig et al., 2018), sugar beet is sensitive to subzero temperatures (Barbier et al., 1982; Loel and Hoffmann, 2015). This sensitivity allows cultivation of the crop in its production area only in a limited vegetation period from spring to late autumn. The limited growth period and the slow formation of leaves in spring are its main yield limiting factors (Jaggard et al., 2009; Milford and Riley, 1980). Calculations taking an increased freezing and bolting tolerance into account suggested that an elongated growth period of sugar beet might result in an increase of the total sugar yield by about 25% (Hoffmann and Kluge-Severin, 2011). However, overwintering sugar beet plants must be able to withstand freezing temperatures, and therefore, freezing tolerance has become a desirable trait for sugar beet breeders. Freezing temperatures frequently lead to severe yield losses of different crops (Barbier et al., 1982; Chang et al., 2014; Fennell, 2004; Maqbool et al., 2010), as intra- and extracellular ice formation damages plant tissue by rupturing cell membranes or because of cellular dehydration (Burke et al., 1976; Wolfe and Bryant, 1999). On the macroscopic level, the impact of freezing on sugar beet taproot tissue is drastic and results in lethal damage to internal tissue and ultimately, the entire plant. Ice formation and thawing leads to cell rupture and leakage of root sap, which ultimately attracts microorganisms and leads to rot of the taproot body (Barbier et al., 1982).

Factors protecting plants against freezing may be soluble sugars, amino acids, organic acids, or derivatives thereof, which function as osmolytes and can thus lower the freezing point, or prevent cellular dehydration upon freezing. Additionally, some of these metabolites can stabilize enzymes, membranes and other cellular components (Guy, 1990; Yadav, 2010). In particular, small carbohydrates like fructans or raffinose family oligosaccharides (RFOs) have superior membrane protective abilities and additionally represent potent quenchers of reactive oxygen species (ROS) (De Roover et al., 2000; Hincha et al., 2007; Pontis, 1989). Accumulation of high levels of fructans in plants of the Asteraceae familiy like chicory (*Cichorium intybus*) or Jerusalem artichoke (*Helianthus tuberosus*), for example, render these species highly tolerant against freezing damage. Sugar beet does not produce fructans but can form considerable amounts of raffinose, which are derived from sucrose, especially during storage of taproots (Haagenson et al., 2008; Wyse and Dexter, 1971). However, raffinose biosynthesis is attended by reduction of sucrose levels. In fact, as part of the so-called molasses, raffinose is considered an unwanted contaminant lowering the maximum amount of sucrose extractable from the pulp during industrial processing (Ganter et al., 1988; Wyse and Dexter, 1971). On the other hand, raffinose is important for plant frost tolerance (Nishizawa et al., 2008; Pennycooke et al., 2003; Peters and Keller, 2009). Raffinose is a trisaccharide consisting of sucrose and galactose and is synthesized via a subsequent transfer of activated galactose moieties, donated by galactinol, to sucrose (Sengupta et al., 2015). Two specific enzymes, galactinol synthase (GOLS) and raffinose synthase (RS), mediate the synthesis of galactinol and raffinose, respectively. GOLS mediates the first metabolic step in raffinose biosynthesis, the conversion of UDP-Galactose and myo-inositol to galactinol. Raffinose biosynthesis is specifically upregulated during cold acclimation processes, and several galactinol synthase genes (*GOLS*) are transcriptionally regulated in *Arabidopsis thaliana* via low temperature response transcription factors of the C-repeat binding factor (CBF) family (Fowler and Thomashow, 2002; Gilmour et al., 2000; Taji et al., 2002). CBF proteins directly regulate cold-responsive (COR) genes, which facilitate the acquisition of cold acclimation and sustained freezing tolerance, as products of COR genes are involved in processes like the synthesis of osmo-protectants and also raffinose, lipid metabolism or cell wall modification (Liu et al., 2019; Thomashow, 1999). In sugar beet, CBFs like CBF3, were shown to be upregulated early upon cold and freezing temperatures, but were only detected in roots so far (Moliterni et al., 2015).

In this work, we analyzed how low temperatures would influence the accumulation of diverse sugar and RFO species in different sugar beet tissues and genotypes. In addition, we present detailed information on raffinose- and inositol-related gene expression and substrate contents in different organs and tissues of sugar beets, exposed to control and freezing temperatures. Our analysis indicates that inositol and raffinose biosynthesis are spatially regulated in a low temperature-dependent manner. We discuss whether these compounds could serve as indicators for the cold and freezing stress level, or whether they may play a direct role in freeze-protection. The identified frost protective substances represent possible targets for screening, breeding or genetic modification of sugar beet varieties with increased winter hardiness and frost tolerance.

## RESULTS

### Sugar beet genotypes respond differently to freezing temperatures

Sub-zero temperatures have a detrimental effect on sugar beet tissue resulting in a structural collapse of the affected cells leading to sucrose leakage out of vacuoles and eventual rotting of taproot tissue (Barbier et al., 1982). To learn about physiological and molecular responses that might mitigate harmful effects of freezing temperatures, we analyzed three genotypes (GT1, GT2, GT3) with different freezing sensitivities from a panel of accessions that was tested for survival of freezing temperatures in field or greenhouse trials. In these tests, GT1 had low survival rates (SR) (14% in field and 11% in greenhouse trials), GT2 high SR (90% in field and 66% in greenhouse trials), and GT3 intermediate SR (77% in field and 56% in greenhouse trials). These rates categorized GT1 as freezing sensitive, GT2 as freezing tolerant, and GT3 as moderately freezing tolerant (**Fig. 1A**). For the experiments presented in this paper, we grew the three different genotypes under a fixed temperature profile in climate-controlled growth chambers and monitored their behavior in respect to the set temperature (**Fig. 1B**). Freezing stress was applied until the temperature, measured 5 cm below the surface of the potting soil, was decreased to 0°C (approximately 3 days). After frost exposure, plants were transferred to 20°C for recovery. The severity of injuries by the three genotypes following treatment indeed mirrored their survival rates from the prior field and greenhouse trials. For instance, most of GT1 shoots collapsed during cold exposure and a comparably high number of GT1 plants did not recover and died after the freezing treatment (**Fig. 1C**). Importantly, initial exposure to freezing temperature did not kill plants, but severely affected their performance during recovery at higher temperatures in a genotype-dependent manner (**Fig. 1C**). Photosynthesis, measured as effective quantum yield of photosystem II (Φ_II_), significantly declined at 0°C in all three genotypes, but strongest in freezing-sensitive GT1 plants (**Fig. 1D**). In contrast, plants from the more freezing-tolerant GT2 and GT3 genotypes were able to increase Φ_II_ to pre-freezing levels after retransfer to 12°C (**Fig. 1D**). Consistently, GT1 plants showed higher non-photochemical quenching (NPQ) in comparison to GT2 and GT3 plants during and after exposure to frost (**Fig. 1D**).

**Figure 1:**
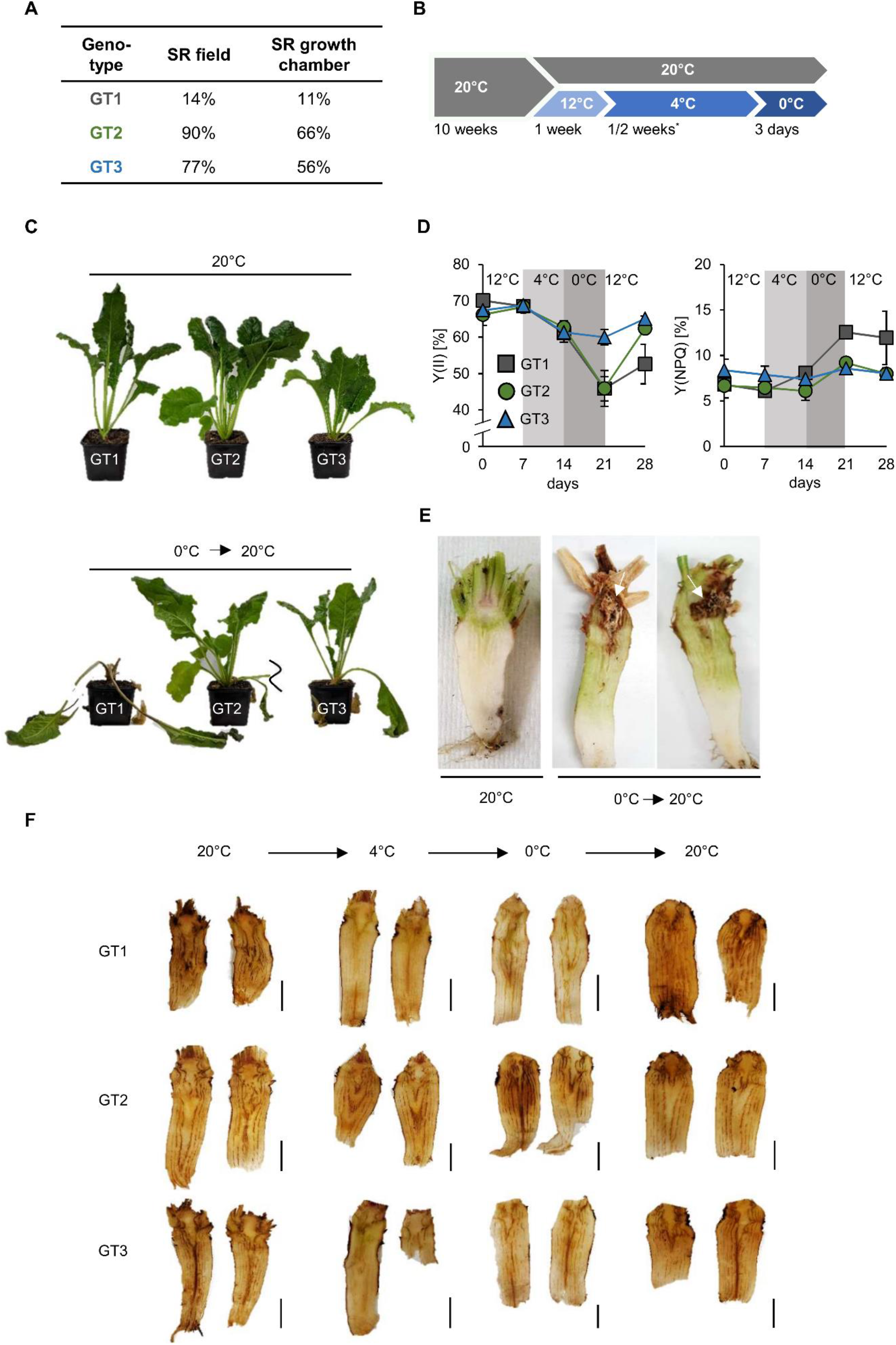
Responses of sugar beet genotypes to freezing temperatures. (A) Freezing survival rates (SR) of three analyzed sugar beet cultivars GT1, GT2 and GT3 measured under field and growth chamber conditions given in percent of survived plants. (B) Experimental setup for metabolite and photosynthesis measurements. Plants were grown at 20°C for 10 weeks. Subsequently, plants were kept at 20°C(control) or transferred to low temperature acclimation conditions (12°C for one week, and 4°C *one week for photosynthesis measurements, two weeks for RNASeq and metabolite analysis). After 4°C treatment, air temperature in the growth chamber was lowered to −6°C and plants were harvested as soon as the soil temperature was 0°C or below, which was the case three days after transfer to −6°C. (C) Typical appearance of sugar beet plants of GT1, GT2 and GT3 before and after freezing treatment. (D) PAM measurements were performed on leaves of the three different genotypes. For this, air temperature was kept at 0°C. Quantum yield of photosynthesis [Y(II)] and non-photochemical quenching [Y(NPQ)] were measured. Temperature intervals are highlighted in white (12°C), light grey (4°C), or grey (0°C). Values represent the mean of three biological replicates. Error bars represent the standard error of the corresponding mean. Significant changes relative to the control condition (20°C) were calculated independently for each genotype using Student’s *t*-test (* p < 0.05; ** p < 0.01; *** p < 0.001). (E) Morphology of sugar beet taproots of genotype GT1 after freezing recovery. Plants were grown at 20°C for 13 weeks and afterwards were shifted to 12°C, 4°C and 0°C and were retransferred to 20°C after freezing treatment. (F) DAB staining indicative for hydrogen peroxide accumulation in longitudinal sections of 10-week old taproots. Scale bars represent 1cm.

Following treatment, a large number of roots from GT1 plants, displayed visible brownish rot in a defined region within the taproot neck, designated as pith (**Fig. 1E**). With progression of the rotting, affected taproots lost structural integrity and plants died. The pith tissues of the two more tolerant cultivars, GT2 and GT3, were not affected to the same extent. Such changes, as observed above, can be evoked by reactive oxygen species (ROS). To check whether the low temperature exposure was associated with formation of ROS, we performed staining of taproot slices with diaminobenzidine (DAB) and nitroblue tetrazolium (NBT) to reveal accumulation of H_2_O_2_ and superoxide, respectively. Staining was most intense during or after freezing treatment and DAB-derived dark brown or NBT-derived blue staining indicated that both types of reactive oxygen species strongly accumulated in the vasculature and in the pith of taproots **(Fig. 1F and Supplemental Fig. S1A)**. Overall, our observations revealed that frost induced tissue damage, as well as emergence of ROS are clustered in the upper part of the sugar beet taproot, mainly in the pith tissue.

### The sugar beet taproot is divided into two distinct tissue zones with different sensitivities to freezing

Almost a century ago, plant anatomist Ernst Artschwager noted the anatomic peculiarities of the pith tissue (Artschwager, 1926). Its anatomy differs from other internal taproot tissues, which mainly comprise the storage parenchyma (SP) and vascular tissue (**Fig. 2A**). Our microscopic analysis confirmed Artschwager’s observation that the pith consists of spongiform parenchyma cells and revealed that pith cells are more than five times the size of parenchymatic cells alternating with xylem/phloem bundles in the distal parts of the beets (i.e. between the characteristic concentric rings that can be observed in transverse sections, mainly through the below-pith beet tissue) (**Fig. 2B-D**). While the storage parenchyma was drawn with vascular bundles, such long-distance transport tissue was absent from the pith, which was surrounded mostly by primary xylem (**Fig. 2C and 2E**). The primary xylem in turn could be traced acropetally towards the vasculature of the oldest leaves and cotyledons, and basipetally to where some of the bundles converge in the central cylinder in below-pith root tissue (**Fig. 2F**). Application of the phloem-mobile carboxyfluoresceine di-acetate (CFDA) to source leaves and subsequent tracking of its green-fluorescent derivative CF in the taproot via confocal microscopy confirmed the occurrence of vascular bundles in storage parenchyma and, importantly, their absence from the pith region (**Fig. 2E and 2F**). To record differential responses in gene expression and metabolite accumulation in SP and pith, we manually separated both tissues. Because of the proximity of the pith to the shoot apical meristem (SAM) **(Fig. 2A)**and with a lack of known markers for the pith, we checked for accumulation of the mRNA of the SAM-marker *CLAVATA2* as indicator for successful enrichment of pith tissue **(Fig. 2G)**. To check whether SP and pith tissue are able to precipitate cold- and freezing treatment under our test conditions, expression of the low temperature response transcription factor *CBF3* was analyzed (**Fig. 2H**). Expression of *CBF3* did not differ between the three analyzed genotypes, but was significantly higher in the pith in comparison to storage parenchyma at 0°C, underlining the marked cold susceptibility of this special tissue (**Fig. 2H**).

**Figure 2:**
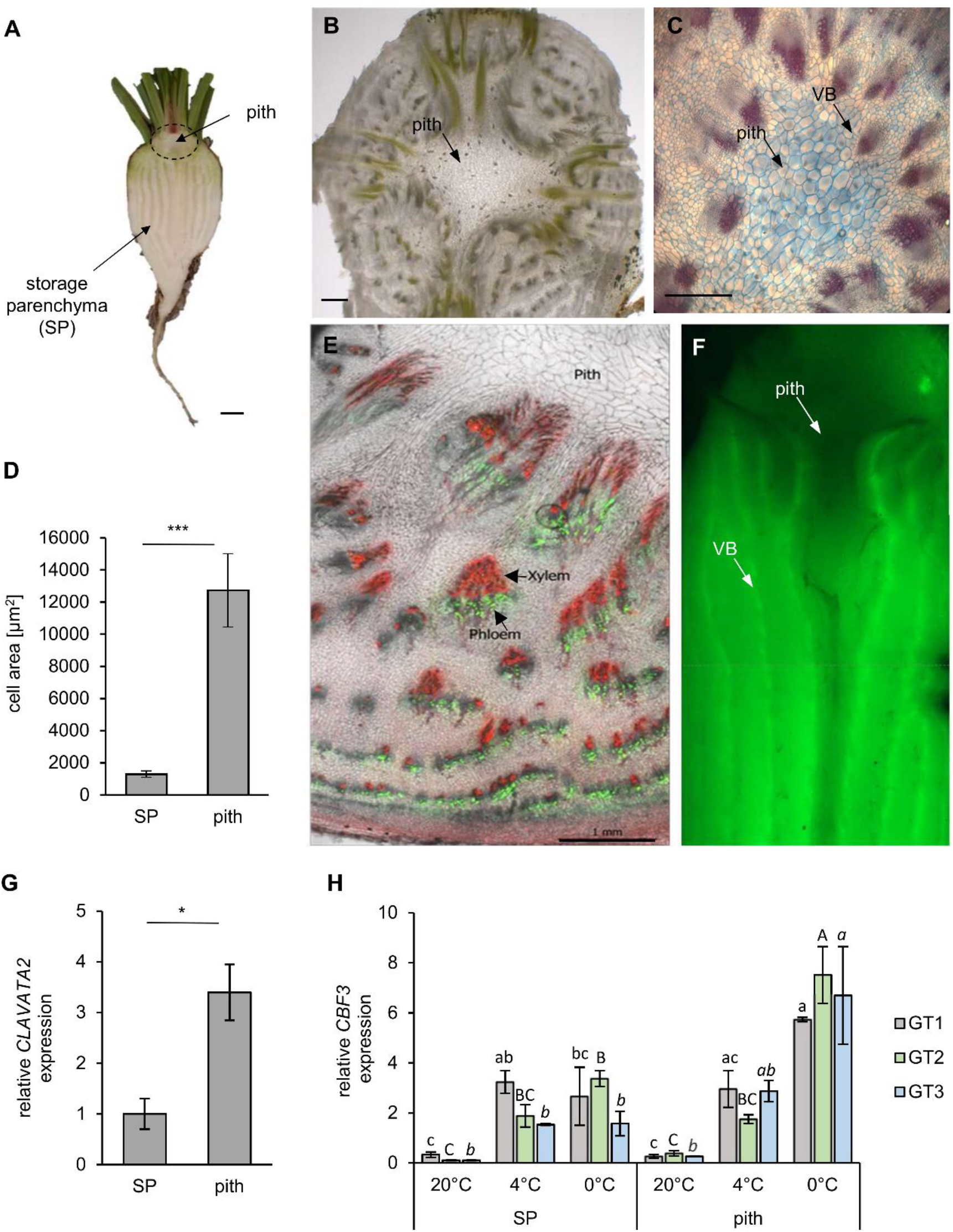
Morphology of the *B. vulgaris* pith tissue. (A) longitudinal overview section of a young taproot. Scale bar representing 1cm. (B) Cross section of the pith tissue to reveal its cellular architecture. Scale bar representing 1 mm. (C) FCA staining of a transverse section of the beet hypocotyl including the central pith. Lignified xylem cells are stained in red, while cellulosic cell walls are stained in blue. Scale bar representing 0.5 mm. (D) Mean cell area of storage parenchyma (SP) and pith cells measured in sugar beet cross-sections using ImageJ. Error bars represent the standard error over seven cross-sections. (E) Radial phloem-unloading of the fluorescent dye carboxyfluorescein (green). Cell walls were stained with propidium iodine (red), which particularly labels the thick, lignified vessels of the xylem. Scale bar represents 1mm. (F) Carboxyfluorescein unloading and distribution from phloem bundles in the sugar beet taproot. Arrows mark the pith tissue or vascular bundles (VB). (G) Pith and storage parenchyma were separated and expression of the shoot meristematic marker *CLAVATA2 was* analyzed. Error bars represent the standard error over three biological replicates. Asterisks indicate significant differences between pith and storage parenchyma (Student’s *t*-test with * p<0.05; *** p<0.001 for (D) and (G)). (H) Expression of the cold-induced transcription factor *CBF3* in SP and pith tissue of the three analyzed genotypes. Error bars represent the standard error over three biological replicates. Letters indicate the same level of significance within each of the three genotypes at different temperatures and in different tissues, calculated via two-way ANOVA with post hoc Tukey test with p< 0.05.

### ROS-related genes are upregulated in the taproot pith at low temperatures

The prominent staining of ROS in freezing-stressed taproots prompted us to analyze antioxidant levels and expression of ROS-related genes in pith and storage parenchyma of the different sugar beet genotypes (**Fig. 3**). Low temperatures lead to a significant increase of the total ascorbate content (sum of ascorbate and dehydro-ascorbate) but did not significantly alter the ratio of oxidized dehydro-ascorbate to reduced ascorbate in the pith tissues of GT1, GT2 and GT3 (**Fig. 3A**). Glutathione also increased at low temperatures reaching highest levels at 4°C in the pith of all three genotypes. The ratio of oxidized GSSG to reduced GSH, serving as an indicator for the ROS stress-level of the tissue, strongly increased with decreasing temperature, reaching its highest value at 0°C in GT1 and GT2, or at 4°C in GT3 (**Fig. 3B**). Compared to the antioxidant contents in the pith, the contents in the storage parenchyma developed similarly with decreasing temperature, but the glutathione contents in the storage parenchyma were overall lower compared to the pith tissue of the different sugar beets (**Supplemental Fig. S1B**). Together with levels of antioxidants, gene expression was monitored in pith and SP tissue during cold- and freezing exposure (**Fig. 3C**). Tissues were analyzed under control temperature (20°C), after one week at 4°C, or after an additional three days at 0°C. Most of the tested genes were upregulated in plants subjected to low temperature treatment, but the increase in expression was always stronger in the pith compared to the SP. Genes encoding typical ROS-scavenging enzymes like catalase, ascorbate peroxidase and reductase, as well as glutathione peroxidase showed the highest expression at 4°C and tuned down again at 0°C. Interestingly, expression of homologs to H_2_O_2_- and superoxide-responsive transcription factor genes *ZAT10* and *ZAT12* increased markedly at 0°C, but not at 4°C (**Fig. 3A**) with highest upregulation observed in the pith tissue, indicating that ROS accumulation increased at 0°C and suggesting that ROS detoxification efficiency was lower at 0°C in comparison to 4°C.

**Figure 3:**
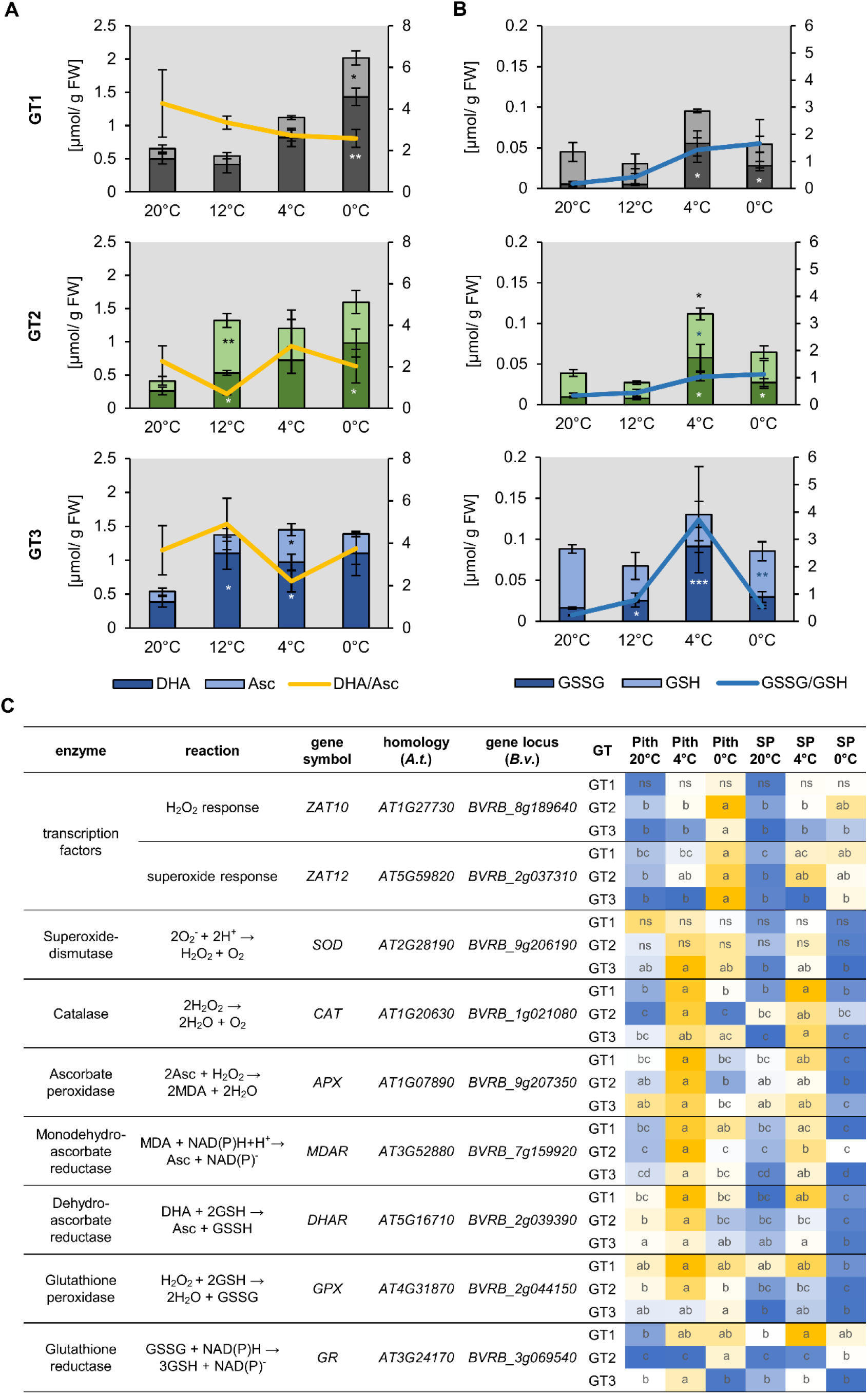
ROS related metabolite accumulation and gene expression. (A) and (B) Concentrations of ascorbate (Asc) and dehydro-ascorbate (DHA) (A) or reduced (GSH) and oxidized glutathione (GSSG) (B) in the pith of different sugar beet genotypes. Reduced form of the antioxidant depicted in light, oxidized form in dark color. Sugar beet plants were grown for 10 weeks at 20°C and were successively transferred to 12°C, 4°C and 0°C for one week each. Three plants of each genotype were harvested at a given temperature and were analyzed for their antioxidant concentration. Error bars represent the standard error over the corresponding mean. Asterisks indicate significant differences in concentrations of reduced, or of oxidized antioxidants or of the oxidized/ reduced ratio calculated in relation to the corresponding values at 20°C using Student’s *t*-test (* p< 0,05; ** p< 0.01). (C) Heat map representation of RT-qPCR analysis of ROS relates genes in taproot pith and storage parenchyma (root) from the three genotypes grown at control (20°C) or low temperatures (4°C and 0°C). Data represents the mean of three independent biological replicates. Different letters within individual tiles denote significant differences between tested conditions and tissues according to two-way ANOVA with post-hoc Tukey HSD testing (p<0.05).

In summary, these results indicate that low temperature-dependent ROS formation and response to ROS accumulation was most pronounced in the pith area of taproots and support the higher accumulation of ROS revealed by ROS staining (**Fig. 1F**).

### The pith tissue is metabolically separated from taproot parenchyma and altered after exposure to subzero temperatures

In addition to the antioxidants ascorbate and glutathione, also other compounds like sugars and amino acids contribute to ROS scavenging. Since ROS accumulation was pronounced in the pith region of taproots, we analyzed the low temperature-dependent accumulation of such metabolites and their spatial distribution between leaf, pith, and storage parenchyma tissues. In total, 36 compounds were measured and their contents are listed in **Supplemental Table S1**. A PC analysis, loaded with the 0°C/20°C fold-changes of these compounds PC1 separated pith and SP tissues (**Supplemental Figure S2**). The metabolic reaction of the storage parenchyma of the two more tolerant genotypes, GT2 and GT3, was similar, but differed from that of GT1 (**Supplemental Figure S2**). The separation of leaf tissue from pith and storage parenchyma was mainly based on changes in raffinose concentrations (**Supplemental Figure S2**). Different degrees of changes in starch content mainly contributed to the separation of pith and storage tissue (**Supplemental Figure S2**). Under control temperatures, the starch content of leaves exceeded that of pith and storage parenchyma about 10-fold (**Supplemental Figure S3**). After exposure to frost, starch decreased drastically in leaves, but was not significantly altered in pith and storage parenchyma (**Supplemental Figure S3**). However, only low concentrations of starch are present in the sugar beet taproot tissue in general and therefore starch is unlikely to contribute significantly to frost-tolerance (Rodrigues et al., 2020; Turesson et al., 2014). Changes in starch content, however, depend on soluble sugar metabolism. Consistently, frost also massively increased concentrations of glucose and fructose in leaves, but not significantly in pith and storage tissues (**Supplemental Figure S3**). Sucrose concentrations were highest in the storage parenchyma and lowest in the leaves (**Supplemental Figure S3**). In both tissues no significant changes in sucrose concentrations could be observed between control and freezing conditions, or between genotypes (**Supplemental Figure S3**). In the pith tissue, concentrations of sucrose differed at 20°C between genotypes, but fluctuated only marginally within and between genotypes at 0°C (**Supplemental Figure S3**).

### Genotype and temperature-dependent accumulation of inositol, galactinol, and raffinose

As concentrations of glucose, fructose and sucrose differed only marginally in pith tissues of the three genotypes after a shift from control to freezing conditions, we analyzed the contents of raffinose as well as of inositol and galactinol, which are precursors for raffinose synthesis (**Fig. 4A**), because raffinose was shown to possess second order rate constants for ROS scavenging, comparable or even higher than those of glucose, fructose and sucrose and the antioxidant ascorbate (Nishizawa et al., 2008). Similar to the sugars glucose, fructose, and sucrose, the pith tissue had inositol, galactinol and raffinose concentrations in-between of those measured for leaves and taproot SP (**Fig. 4**). Inositol content of the leaves at 20°C was highest in GT2 and GT3. Upon 0°C exposure, inositol levels increased to comparable amounts in the leaf tissue of all three genotypes (**Fig. 4B**). However, in the pith and storage parenchyma, inositol levels remained unchanged upon exposure to freezing temperatures (**Fig. 4B**). Like its precursor inositol (**Fig. 4A**), concentrations of galactinol increased in leaves at 0°C two-to three-fold **(Fig. 4C)**. Galactinol concentrations also increased in the pith and storage tissue at 0°C. In the pith galactinol accumulated at 0°C to concentrations twice as high as compared to the SP (**Fig. 4C**). Notably, freezing temperatures boosted raffinose concentrations at least 10-fold, in GT3 leaves, and up to 80-fold, in GT1 leaves, in comparison to the corresponding leaf-tissue at 20°C (**Fig. 4D**). In the pith tissue, raffinose content increased in GT2 and GT3 at 0°C. At 0°C, raffinose levels of GT2 and GT3 exceeded those of GT1 at least two-fold in the pith. Interestingly, GT2, the cultivar with the highest freezing survival rates (**Fig. 1A**), had about 80% higher raffinose concentrations in comparison to GT1 and GT3 already under control conditions (**Fig. 4D**). In SP, raffinose concentrations were slightly increased or remained unchanged at 0°C in comparison to 20°C (**Fig. 4D**).

**Figure 4:**
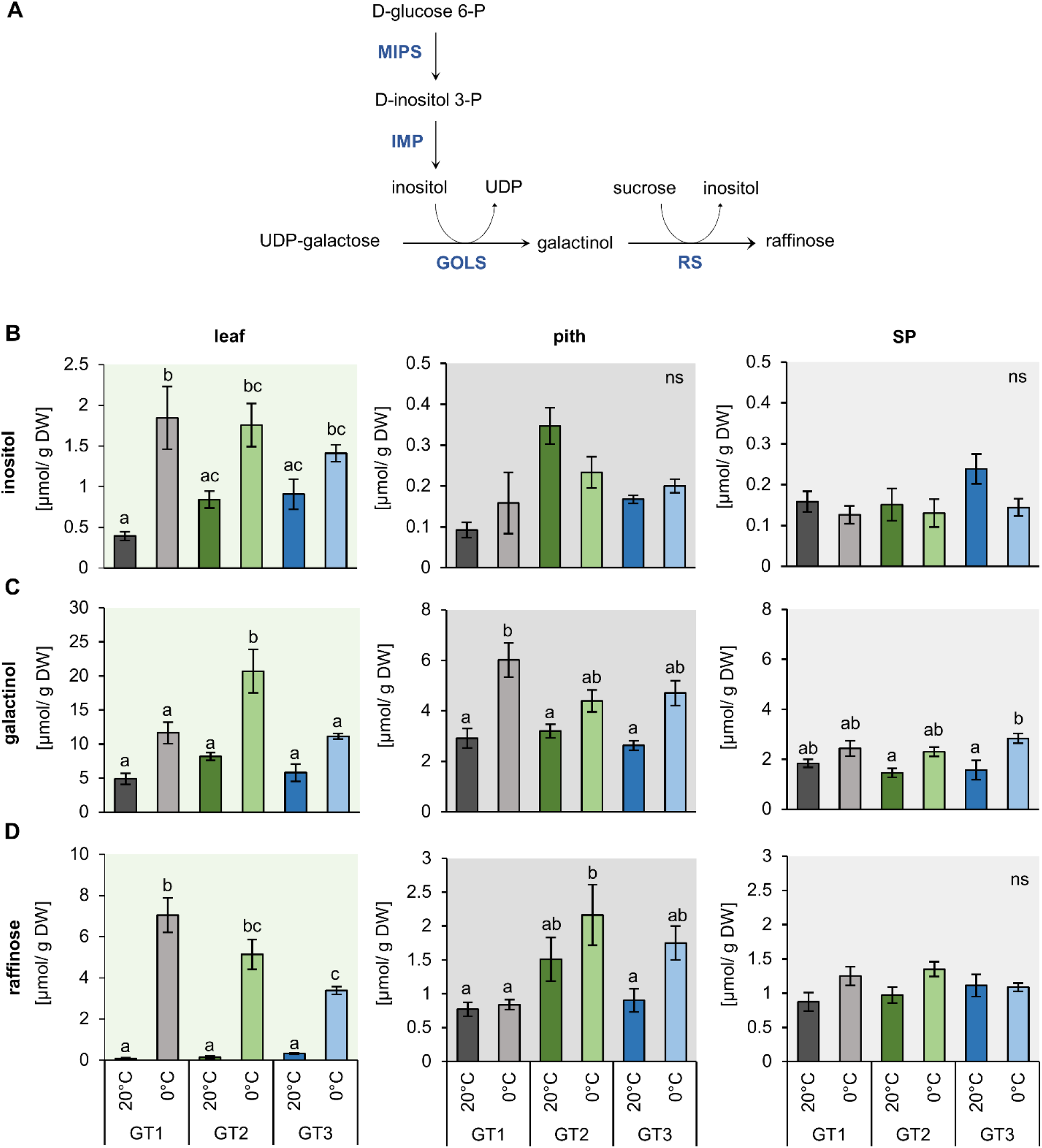
Inositol and raffinose synthesis and their concentrations in leaf, pith and storage parenchyma tissue under control and freezing temperatures. (A) Schematic representation of inositol and raffinose synthesis in plants. (B) to (D) Raffinose and intermediates in raffinose synthesis were measured in plants grown at 20°C and plants transferred to 12°C, 4°C and harvested at 0°C soil temperature. Plants were dissected in the three different tissues leaf, pith and root while harvesting at the corresponding temperatures. Inositol (B), galactinol (C) and raffinose (D) concentrations represent the mean of three biological replications for each of the tested cultivars. Error bars represent the standard error over the corresponding mean. Letters indicate the same level of significance for each measured concentration, calculated via two-way ANOVA with post hoc Bonferroni test with p< 0.05.

### Inositol-, galactinol- and raffinose-biosynthesis genes are differentially expressed in leaf and taproot tissue in response to freezing

To reveal global patterns of gene expression in response to freezing stress we performed comparative RNASeq analysis of leaf and taproot tissues of the three different genotypes at control and freezing temperatures. However, this analysis came with the limitation that we did not separate pith and storage parenchyma but instead analyzed total taproots. RNA-Seq reads were mapped to the sugar beet reference genome (Dohm et al., 2013) and revealed global rearrangement of gene expression in leaf and taproot tissue upon freezing exposure (**Fig. 5A, B and Fig. 5F, G**).

**Figure 5:**
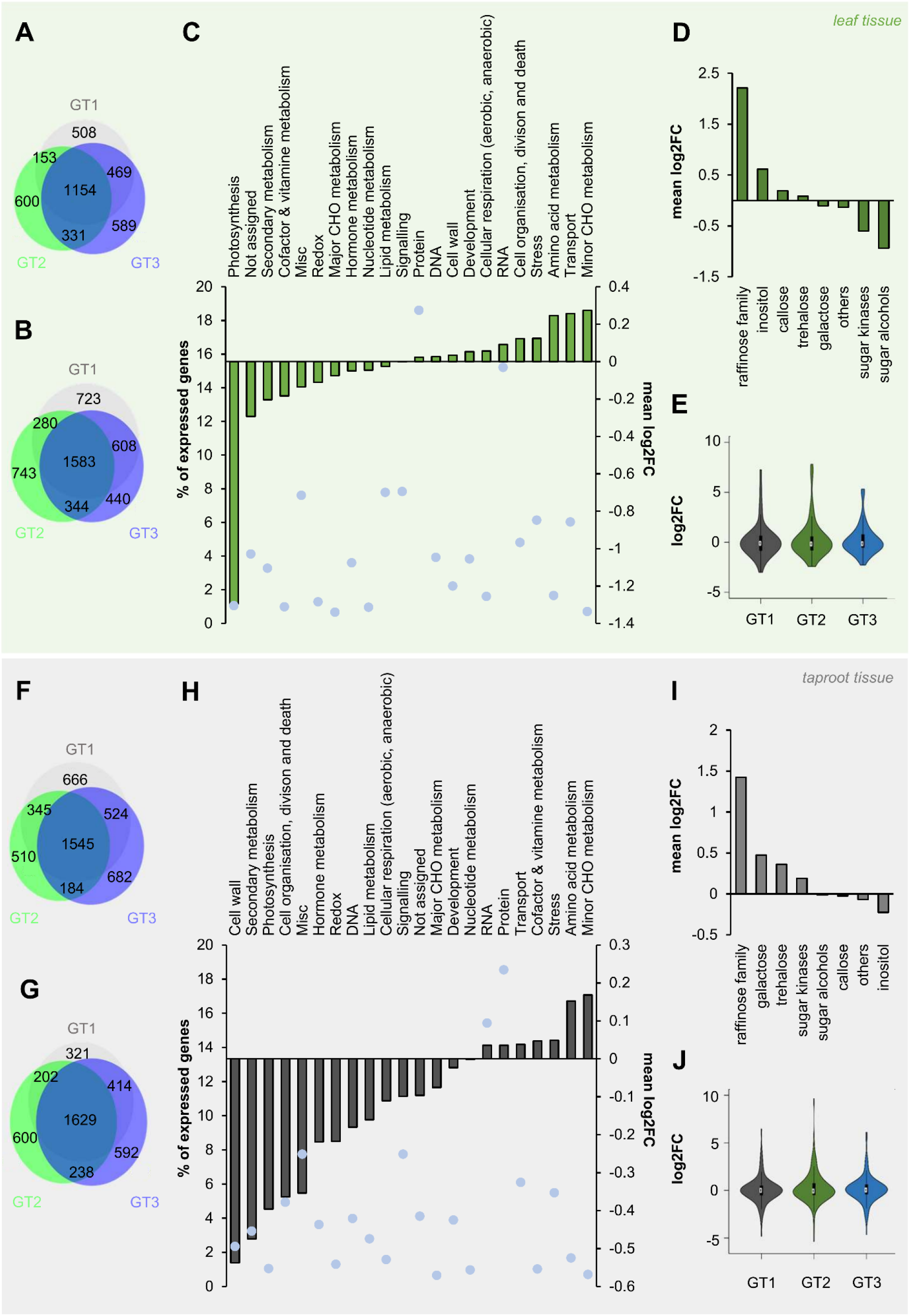
Changes in gene expression of *B. vulgaris* leaf and taproot tissues upon freezing treatment. (A, b; F, G) Venn diagrams of differentially expressed genes in leaves (A, B) and taproots (F, G). Numbers of up – (Log2FC ≥ 1; A, F) or down – (Log2FC ≤ −1; B, G) regulated genes (with a FDR ≤ 0.05) in leaves (A, B) or taproots (F, G) are given inside the circles of the diagram. Log2FC 20°C/ 0°C of all expressed genes with log2FC ≠ 0 in leaf (C) and taproot tissue (H) were functionally grouped and plotted against the number of genes in each group given in percent of the total number of expressed genes (blue dots). (D) and (I) mean log2FC 20°C/ 0°C of genes with log2FC ≠ 0 of different metabolic pathways included in the functional group of “minor CHO metabolism” in leaf (D) and taproot tissue (I). Log2FCs represent the mean of the three different *B. vulgaris* genotypes GT1, GT2 and GT3, each measured in three biological replications for (C, D) in the leaf and (H, I) in the taproot. Log2FCs of genes included in the functional group of “minor CHO metabolism” are shown as violin plots for the corresponding tissues leaf (E) and taproot (J). White circles show the medians, box limits indicate the 25th and 75th percentiles as determined by R software, whiskers extend 1.5 times the interquartile range from the 25th and 75th percentiles and polygons represent density estimates of data and extend to extreme values in violin plots.

To get a more general overview on regulation of different cellular pathways and reactions, RNASeq data were mapped into different functional groups, based on MapMan mapping. We then calculated the mean logarithmic fold change (log2FC) of all genes with log2FC ≠ 0 mapped in each of these functional groups, without filtering for significant changes prior to analysis. This analysis revealed downregulation of genes involved in photosynthesis in leaf tissue at 0°C (**Fig. 5C**). In contrast, minor CHO metabolism was the most prominently upregulated functional group in the leaf (**Fig. 5C**) and also in the taproot tissue (**Fig 5H**). Minor CHO metabolism separates into the sub-bins raffinose, inositol, sugar alcohols, galactose, trehalose, callose synthesis, and sugar kinases. The mean log2FCs of genes involved in each of these sub-bins ranged from lowest values for sugar alcohols to highest for raffinose metabolism in leaves and from lowest mean log2FC for inositol to highest for raffinose metabolism in taproot tissue (**Fig. 5D and 5H**). The latter opposite regulation of inositol and raffinose in taproots is interesting because inositol is essential for raffinose biosynthesis and the high log2FCs for raffinose in both leaves and taproots suggested increased utilization of carbon for raffinose biosynthesis under low temperatures. To reveal first genotype dependent differences in the expression of minor CHO genes in the different genotypes, violin plots showed the distribution of log2FC over minor CHO genes in leaf and taproot tissues. While most of those genes show log2FCs between −2 and 2 in both leaf and taproot tissues upon freezing treatment (**Fig. 5E and 5J**), in GT2 a few genes showed log2FC of up to 7 in the leaves and up to 10 in the taproot (**Fig. 5E and 5J**). Overall the number of highly upregulated genes in minor CHO metabolism of GT2 was higher than in GT1 and GT3, independent of the tissue (**Fig. 5E and 5J**). To reveal possible genotype specific regulation of genes involved in raffinose and its precursors synthesis, we extracted transcript levels for those genes upon exposure to 0°C (**Table 1**).

**Table 1:**
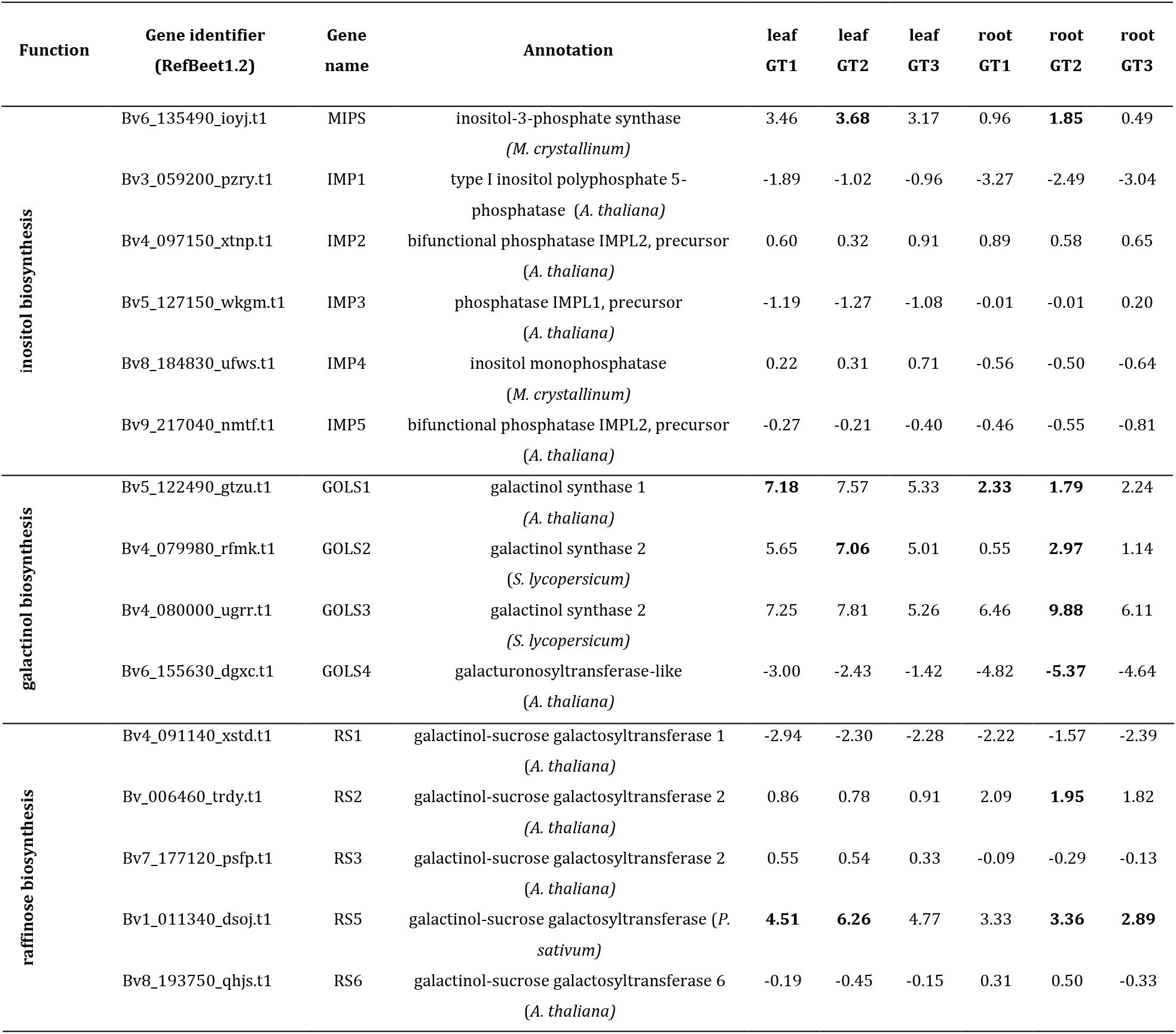
Log2FC 20°C/ 0°C of expression of enzymes involved in inositol-, galactinol and raffinose synthesis of leaf and taproot tissue. (root) of three differential *B. vulgaris* genotypes GT1, GT2 and GT3. Data have been extracted from the RNASeq analyis. Significant up- or downregulation at 0°C in comparison to the expression at 20°C is indicted by bold numbers. Significance was calculated using Student’s *t*-test with false discovery rate testing (padj <0.1).

*De-novo* inositol biosynthesis occurs via the enzymes myo-inositol phosphate synthase (MIPS) and inositol monophosphatase (IMP) (**Fig. 4A**; Loewus and Murthy, 2000). MIPS catalyzes the rate limiting step in inositol *de-novo* synthesis (Donahue et al., 2010), isomerization of glucose-6-phosphate to inositol-3-phosphate. IMP then catalyzes de-phosphorylation of inositol-3-phosphate to inositol. In the *B. vulgaris* genome (Dohm et al., 2013), we identified one sequence with high homology to MIPS (*Bv*MIPS, Bv6_135490_ioyj.t1) and five sequences with high homology to IMP (*Bv*IMP1, Bv3_059200_pzry.t1; *Bv*IMP2, Bv4_097150_xtnp.t1; *Bv*IMP3, Bv5_127150_wkgm.t1; *Bv*IMP4, Bv8_184830_ufws.t1; *Bv*IMP5, Bv9_217040_nmtf.t1).

Frost enhanced expression of *BvMIPS* in both, leaves and taproots, with the strongest induction occurring in the freezing-tolerant cultivar, GT2 (**Table 1**). Expression of the five *IMP* genes was differentially regulated upon 0°C. In leaves, *IMP1, IMP3*, and *IMP5* were downregulated, while *IMP2* and *IMP4* genes were slightly upregulated at 0°C. In taproots, IMP expression was downregulated, except for IMP2 (**Table 1**). However, *IMP* genes were not regulated in a genotype-dependent manner.

Inositol can be fused to UDP-galactose to form alpha-D-galactosyl-(1->3)-1D-myo-inositol (galactinol) and UDP. This reaction is catalyzed by galactinol synthase (GOLS) and is required for subsequent biosynthesis of raffinose (**Fig. 4A**). We identified four genes with homology to galactinol synthases from tomato (*Solanum lycopersicum*) or Arabidopsis in the sugar beet genome (**Table 1, Supplemental Figure S4**). The expression of three of the four *GOLSs* (*BvGOLS1 to 3*) was strongly upregulated at 0°C in leaves and in taproots. The log2FCs for *GOLS2* and *GOLS3* in leaves and especially in the taproot of GT2 exceeded those of the less freezing tolerant genotypes GT1 and GT3 (**Table 1**). The gene coding for a fourth GOLS isoform, *GOLS4*, was downregulated at 0°C, independent of the tissue and genotype (**Table 1**). Phylogenetic analysis revealed that *BvGOLS4* (Bv6_155630_dgxc.t1) was only distantly related to the other sugar beet GOLS isoforms (**Supplemental Figure S4**) and therefore may not encode a functional GOLS or a GOLS relevant for the plants freezing response.

The ultimate step of raffinose biosynthesis is the transfer of the galactinol’s galactose moiety to sucrose via the enzyme raffinose synthase (RS). During this reaction, inositol is released from galactinol (**Fig. 4A**). We identified seven RS isoforms in the sugar beet genome (**Supplemental Figure S4**). Two isoforms, *BvRS4* and *BvRS7* were not changed in their expression. Of the remaining 5 isoforms, two RS, *BvRS2* and *BvRS5*, were upregulated in the leaf and taproot tissue at 0°C (**Table 1**). Overall, *BvRS5* (*Bv1_011340_dsoj.t1*), the closest homolog to *AtRS5* (**Supplemental Figure S4**), showed the highest upregulation at 0°C (**Table 1**). Similar to the GOLS genes, upregulation was strongest in freezing-tolerant GT2 plants, independent of the tissue (**Table 1**).

### Raffinose biosynthesis genes are induced in pith tissues at low temperatures in a genotype-dependent manner

The RNAseq analysis suggested that raffinose metabolism is important in both leaf and taproot tissues during the freezing response of sugar beet plants (**Figure 5 and Table 1**). Because of the different sensitivities of the pith of the three genotypes to freezing, we analyzed the expression of galactinol and raffinose biosynthesis-related genes in taproot tissues with RT-qPCR (**Fig. 6**). Among galactinol synthases, *GOLS1* was the most abundant isoform in the pith and storage parenchyma (**Fig. 6A**), however, upregulation of gene expression in terms of fold-change was strongest for *GOLS2* and *GOLS3* at 0°C (**Fig. 6A**). Expression of *GOLS2* and *GOLS3* was highest in GT1 and GT3 at 0°C (**Fig. 6A**). The two RS isoforms *RS2* and *RS5* showed comparable abundance in the pith of all three genotypes at control temperatures. Abundancy of *RS2* was slightly higher in the storage parenchyma, while *RS5* abundancy in the SP was comparable to the pith at 20°C (**Fig. 6B**). Additionally, the upregulation of RS expressions in terms of fold-change in the pith was comparable between *RS2* and *RS5* expression in GT2 and GT3. Overall, GT3 showed the highest fold-change upregulation of *RS2* and *RS5* in the pith (**Fig. 6B**). In GT1 pith tissue, upregulation of *RS2* expression was comparable to that in GT2 (**Fig. 6B**). While *RS2* was upregulated in the pith of GT1 upon exposure to 0°C, GT1 pith tissue lacked the ability of upregulation of *RS5* upon exposure to freezing temperatures (**Fig. 6B**). While *RS5* expression increased more than 15-, and 20-fold in the pith of GT2 and GT3 respectively, expression remained unaltered in the pith of the less tolerant GT1 upon 0°C (**Fig. 6B**). GT1 failed to induce *RS5* in the pith tissue in comparison to GT2 and GT3 and induction of *RS5* expression was highest in SP of this genotype (**Fig. 6B**). Taken together, expression of *RS* genes in the pith was upregulated to a greater extent in the freezing-tolerant cultivars GT2 and GT3, and to a lower extent in sensitive GT1 (**Fig. 6B**), which is in line with the low raffinose concentrations in the pith of this genotype at 0°C (**Fig. 4D**).

**Figure 6:**
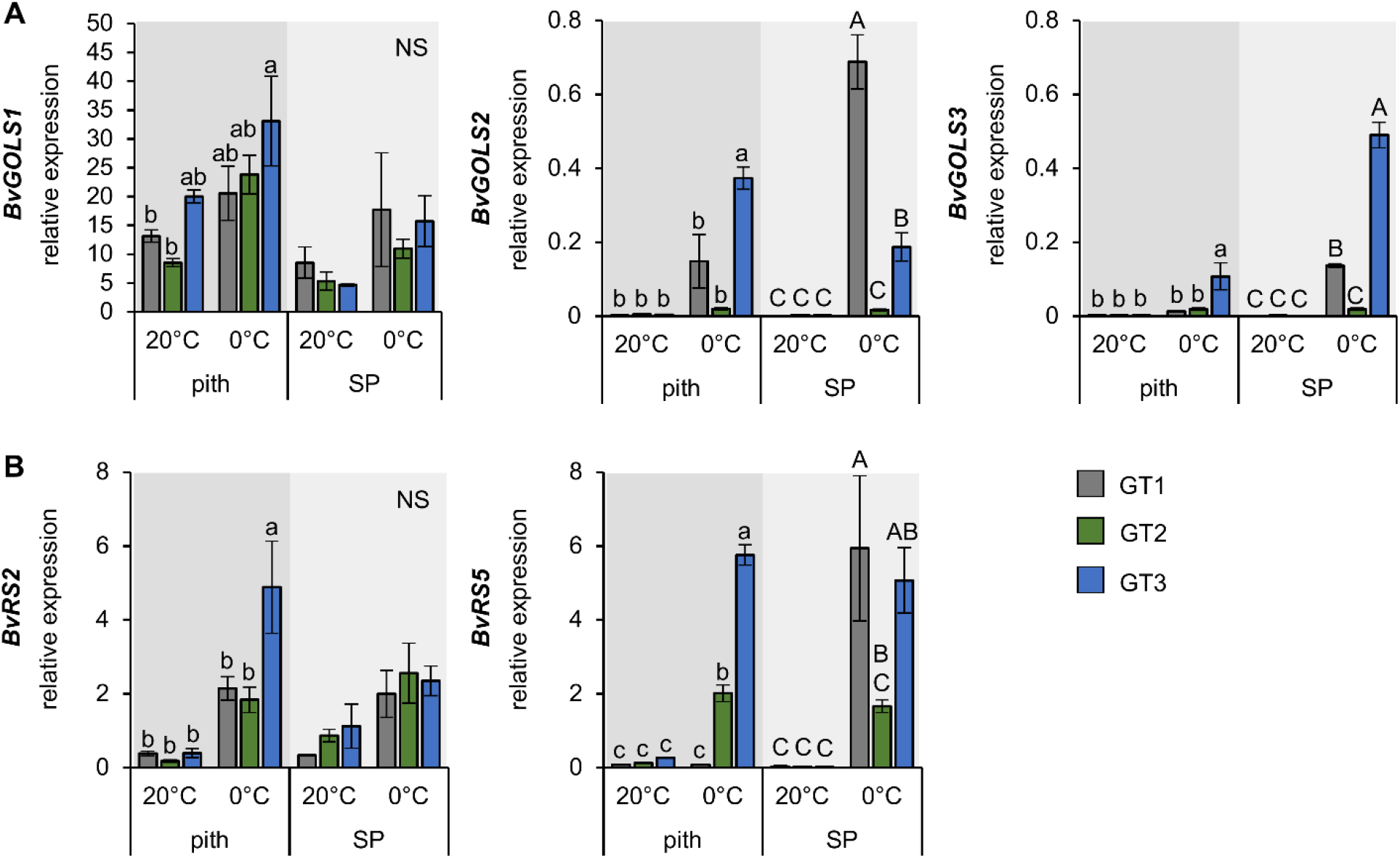
Expression of *GOLS* and *RS* isoforms in pith and storage parenchyma samples of three contrasting *B. vulgaris* cultivars. Relative expression of *GOLS* (A) and *RS* isoforms (B) in the pith and storage parenchyma at 20°C and freezing treatment at 0°C. Bars represent means from three biological replicates ± standard error. Different letters indicate significant differences between genotypes within a tissue according to two-way ANOVA with post hoc Tukey test (p< 0.05).

## DISCUSSION

During exposure to low temperatures (4°C), sugar beet plants undergo a transformation in metabolite- and gene expression profiles (Rodrigues et al., 2020). However, as we show here, not only chilling but also freezing treatment induces changes in metabolite concentrations and gene expression. Because *Beta vulgaris* is a freezing-sensitive plant showing severe freezing-induced tissue damage (**Fig. 1C**), it is of high interest to analyze the plant’s response to sub-zero temperatures on a molecular basis.

Transcriptomic analysis revealed that genes involved in minor CHO metabolism were strongly upregulated in both leaf- and taproot tissues (**Fig. 5 and Table 1**). While genes from this functional group were upregulated, genes involved in photosynthesis were downregulated upon freezing treatment (**Fig. 5**). Both the downregulation of photosynthesis-related genes and the decrease in photosynthetic activity (**Fig. 1D**) at low temperatures confirmed our recent results, where cold (4°C) treatment led to lower photosynthetic activity and downregulation of photosynthesis-genes in sugar beet leaves (Rodrigues et al., 2020). Such effects on photosynthetic related protein level has also been observed in leaves of Arabidopsis (Fanucchi et al., 2012; Janmohammadi et al., 2015) and may be a direct effect of electron overflow at photosynthetic reaction centers because of a slow-down of Calvin-Benson cycle reactions (Pommerrenig et al., 2018; Strand et al., 1997).

Sugar concentrations, especially those of monosaccharides, increased at 0°C in the leaf tissue (**Supplemental Figure S3**) in a similar manner as in other plant species, where metabolites accumulate during prolonged cold treatment (Ho et al., 2020; Klemens et al., 2014; Usadel et al., 2008). Overall, concentrations of sucrose were highest in the heterotrophic tissues, whereas concentrations of the monosaccharides glucose and fructose were highest in the leaves (**Supplemental Figure S3**). This pattern was expected, as sucrose is the main storage carbohydrate of taproots (Giaquinta, 1979; Rodrigues et al., 2021). The pith tissue harbored intermediate concentrations of both, monosaccharides and sucrose, as - in contrast to the storage parenchyma – it is not directly supplied with sucrose by the phloem. The application of the phloem mobile stain CFDA (Knoblauch et al., 2015; Mehdi et al., 2019; Pommerrenig et al., 2020) to source leaves allowed visualization of phloem fluxes into and within the sugar beet taproot and indicated that sucrose was transported in the taproot via the phloem in a similar manner. CFDA staining revealed organization of the phloem in vascular rings surrounding the pith (**Fig. 2**). Sugar translocated via and unloaded from the phloem can therefore not directly enter the pith tissue (**Fig. 2**) (Artschwager, 1926; Jahnke et al., 2009; Metzner et al., 2014). This observation correlates with lower amounts of sucrose in the pith tissue (**Supplemental Figure S3**). The size of pith cells was markedly larger in comparison to storage parenchyma cells and morphology of the pith already suggests lower sucrose concentrations in its cells, since sucrose concentration has been reported to decline with cell size in sugar beet (Bellin et al., 2007).

Our observations show that post-freezing rot of taproot tissue mainly originates from the pith tissue. High ROS accumulation at freezing temperatures as indicted by intense DAB- and NBT-derived staining (**Fig. 1 and Supplementary Figure S1**) and strong upregulation of ROS marker genes *ZAT10* and *ZAT12* (**Fig. 3**) might be causative for the observed tissue damage resulting in post-freezing rotting of the pith. Genes encoding factors of the ROS detoxification system were markedly upregulated at chilling temperatures (4°) or freezing temperatures (**Fig. 3**). Raffinose biosynthesis genes on the other hand were also increased at 4°C but maintained high mRNA levels at 0°C (**Fig, 5 and Table 1**).

Raffinose is known to accumulate under abiotic stress conditions such as drought (Li et al., 2020) or cold in different plant species including Arabidopsis and rice (*Oryza sativa*) (Saito and Yoshida, 2011; Taji et al., 2002). Raffinose is synthesized by a transfer reaction of galactosyl units to sucrose, involving the enzymes galactinol synthase (GOLS) and raffinose synthase (RS) (**Fig. 4A**). Both enzymes were reported to be upregulated during cold treatment in *Ajuga reptans, Medicago falcata, Medicago sativa* or *Arabidopsis thaliana* (Cunningham et al., 2003; Egert et al., 2013; Sprenger and Keller, 2000; Zhuo et al., 2013; Zuther et al., 2004). The enzymes myo-inositol phosphate synthase (MIPS) and inositol monophosphatase (IMP) are indirectly involved in raffinose synthesis, as they catalyze the *de novo* synthesis of inositol, which is required for the formation of the galactosyl donor galactinol (Saravitz et al., 1987; Sengupta et al., 2015). *MIPS* and *IMP* genes were also shown to be upregulated at low temperatures in plants like rice (*Oryza sativa*), yellow passion fruit (*Passiflora ligularis*), or chickpea (*Cicer arietinum*) and have been implicated to play a role in environmental stress responses by promoting RFO biosynthesis (Abreu and Aragão, 2007; Kaur et al., 2008; Saxena et al., 2013; R.-X. Zhang et al., 2017).

We observed that raffinose synthesis was highly upregulated upon freezing temperatures (**Fig. 5 and Table 1**) suggesting a central role for the involved enzymes in sugar beet freezing response. Among the four genes coding for GOLS isoforms in the sugar beet genome (**Table1 and Supplemental Figure S5**), three were upregulated upon exposure to freezing, including the two closest homologs to GOLS2 and GOLS3 from Arabidopsis. (**Table 1 and Supplemental Figure S4**). *AtGOLS2* and *AtGOLS3* are both known to be upregulated specifically by drought and cold stress (Maruyama et al., 2009; Shen et al., 2020; Taji et al., 2002) and their induction involves DREB1A/CBF3 transcription factors (Maruyama et al., 2009). These transcription factors are induced during exposure to low temperatures (Gilmour et al., 1998; Shinwari et al., 1998; **Fig. 2H**), and their expression has been shown to confer freezing tolerance in Arabidopsis, rice, or perennial ryegrass (*Lolium perenne*) (Gilmour et al., 2000; Ito et al., 2006; Li et al., 2011).

Seven raffinose synthase isoforms were identified in sugar beet, of which two were not differentially regulated upon exposure to low temperatures and three isoforms were differentially regulated in leaf and taproot tissues (**Table1 and Supplemental Figure S4**). The two close homologs of *AtRS2* and *AtRS5* were upregulated at freezing temperatures and showed the most pronounced induction in the freezing tolerant cultivars GT2 and GT3 (**Table 1**). While AtRS2 was shown to possess α-galactosidase activity (Peters et al., 2010), AtRS5 was proven to be a raffinose synthase, upregulated in the cold (Egert et al., 2013). Interestingly, *BvRS5*, the closest homolog to AtRS5, was also highly upregulated in the piths of the two cultivars with higher freezing tolerance, GT2 and GT3, but not in freezing-sensitive GT1 plants (**Fig. 6B**).

Upregulation of *MIPS*, *IMP*, *GOLS* and *RS* isoforms went along with high accumulation of inositol, galactinol and raffinose upon exposure to freezing temperatures (**Fig. 4**). Interestingly, raffinose levels in the pith of more cold tolerant genotypes differed from concentrations in the pith of the susceptible genotype – even in the absence of cold (**Fig. 4D**). GT2, the cultivar with the highest freezing tolerance (**Fig. 1A**) and also the highest upregulation of several *GOLS* and *RS* isoforms (**Table 1**) shows the highest raffinose amounts in the pith region amongst the three genotypes, already at 20°C. In GT3, increase in raffinose concentration upon subzero temperatures is highest, most likely mediated by the high upregulation of *RS5* specifically in the pith tissue (**Fig. 6B and Fig. 4D**). In GT1, raffinose concentrations were not altered by freezing temperatures, which is in line with the expression of raffinose synthases, which were not induced in the pith (**Fig. 6B and Fig. 4D**). These data suggest that survival of the pith tissue might be linked to high raffinose concentrations and are in line with previous reports, that showed that raffinose levels were elevated upon cold and freezing temperatures (Koehler et al., 2015; Yue et al., 2015). Such conditions induce the formation of reactive oxygen species (Gechev et al., 2006), which, when accumulating to high levels, can be harmful to the plant, as they can damage membranes (lipid peroxidation), proteins, RNA and DNA molecules (Choudhury et al., 2017). Accumulation of galactinol and raffinose on the other hand, can diminish those harmful effects of ROS. This is because, high levels of these metabolites were shown to stabilize ascorbate and glutathione concentrations, which are involved in detoxification of ROS concentrations (Keller et al., 2021; Nishizawa-Yokoi et al., 2008). Furthermore, galactinol and raffinose show second-order rate constants for detoxification of hydroxyl radicals higher than those of common antioxidants like ascorbate or glutathione (Nishizawa-Yokoi et al., 2008). Although raffinose is synthesized in the cytosol, about 20% of cellular raffinose is located to the chloroplasts and its concentration in plastids is increased even further at low temperatures (Knaupp et al., 2011; Schneider and Keller, 2009). The majority of cellular raffinose is, however, located in the vacuole, reaching about 60% in cold acclimated Arabidopsis plants (Knaupp et al., 2011). In comparison to leaf tissue, taproot tissue does hardly contain any chloroplasts and the vacuole is most likely the major place for raffinose accumulation under low temperatures. There, raffinose might accumulate near the tonoplast together with other carbohydrates. Excess cytosolic H_2_O_2_ that passes through the tonoplast, or H_2_O_2_ originating from superoxide produced by tonoplast-localized NADPH oxidases, could then react with raffinose, similar to the reaction proposed for fructans (Matros et al., 2015; Peshev et al., 2013), to protect the tonoplast from membrane damage via ROS. In addition to its important role in ROS detoxification, RFOs can assist in the osmotic adjustment and thereby maintain cell turgor during water loss due to freezing (ElSayed et al., 2014) and protect the cell against severe damage. The higher raffinose concentrations of the freezing tolerant cultivars GT2 and GT3 in comparison to freezing sensitive GT1, especially in the freezing damage prone pith tissue, hint towards a beneficial contribution of raffinose for freezing protection of this delicate tissue.

In Arabidopsis, ICE-CBF/DREB1-regulation is the main low temperature-responsive pathway leading to raffinose accumulation (Janmohammadi et al., 2015) and in our experiments sugar beet *CBF3* was highly expressed at freezing temperature in the pith (**Fig. 2H**). It is tempting to speculate that the regulation of the same pathway might cause an upregulation of specific GOLS and RS isoforms also here in *B. vulgaris*, as the closest sugar beet homologs to *AtGOLS2*, *AtGOLS3* and *AtRS5* all show similarly high upregulation upon freezing temperatures (**Table 1, Supplemental Figure S4**). This suggests that accumulation of galactinol and especially of raffinose is a regulated acclimation reaction of sugar beet plants rather than a result of an altered metabolism in the context of a general stress response.

## MATERIAL AND METHODS

### Plant Material and Growth conditions

Three hybrid sugar beet genotypes with contrasting winter-hardiness (GT1, GT2, GT3; KWS SAAT SE, Germany) were used. Plants were germinated and grown on standard soil substrate ED73 (Einheitserdwerke Patzer, Germany) with 10% (v/v) sand under a 10 h light/14 h dark cycle, 60% relative humidity, and 110 μmol m^−2^ s^−1^ light intensity. Plants were grown at 20°C for 10 weeks. The plant population was split and one group was transferred to 12°C for one week and 4°C for one to two weeks (one week for ROS and Photosynthesis measurements; two weeks for metabolite measurements and gene expression) afterwards. After 4°C treatment, plants were transferred to −6°C air temperature and were kept there until soil temperature reached 0°C, which took approximately three days (**Figure 1B**). Control plants were kept at 20°C. During harvest, plants were dissected into leaf, pith and storage root tissues, using a kitchen knife. The pith tissue was identified visually from beets halved lengthwise by its peculiar cellular morphology (**Figure 1E and Figure 2**) by eye. Tissue from two different plants was pooled and treated as one independent replicate, with a total of four replicates per genotype and condition, respectively. After harvest, plant material was transferred to liquid nitrogen. Samples used for metabolite analysis were lyophilized in an Alpha 2-4 LD plus freeze-drier (Christ, Osterode am Harz, Germany) for seven days. Sampled material was pulverized using the Retsch MM301 mill (Retsch, Haan, Germany) with a frequency of 30 s^−1^ for 30 seconds.

### Survival rate testing

For survival rate testing of different sugar beet genotypes, greenhouse and field experiments were carried out. Therefore, plants were sown out approximately three months before start of the frost period. During experiments it was made sure, that plants were subjected to soil temperatures below 0°C. At least two weeks after frost treatment, dead or rotten sugar beets plants were counted and survival rate was calculated. The survival rates monitor mean values of at least two replications in independent years and at different locations.

### Chlorophyll Fluorescence Measurements

Photosynthetic activity of three individual plants per genotype was measured using an Imaging-PAM *M-Series-System* (Heinz Walz, Effeltrich, Germany). Plants were placed in the dark for 8 min to deplete the energy of PSII. Capacity of PSII was measured by saturation with 14 cycles of photosynthetically active radiation (76 μmol photons m^−2^ s^−1^) light-pulses at 0, 50, and 70 s. Recorded fluorescence was used for calculation of effective quantum yield of PSII (Y(II)) and quantum yield of nonphotochemical quenching (Y(NPQ)).

### RNASeq analysis

For RNASeq analysis plants were grown according to the growth regime mentioned by Martins Rodrigues et al. 2020. Additionally, RNA extraction and analysis of RNASeq results including statistical analysis was performed as described in Martins Rodrigues et al., 2020 and data was made publicly available as BioProject PRJNA602804. For general overview of RNASeq results annotations were assigned to the corresponding MapMan bincode found in the mapping file named “Ath_AFFY_STv1.1_TRANSCRIPT_CLUSTER_TAIR10_LOCUS”. Means over the log2FC over the three independent genotypes were calculated for each bincode for leaf and taproot tissue, containing every log2FC ≠ 0 and without filtering for significant changes.

### Expression analysis via RT-qPCR

Expression analysis of selected genes was performed by reverse transcription quantitative PCR (RT-qPCR) using the primers listed in **Supplemental Table S2**. RNA was extracted from homogenized sugar beet tissue using the NucleoSpin(R) RNA Plant Kit (Machery-Nagel, Düren, Germany) according to the manufacturer’s guidelines. RNA transcription into cDNA was performed using the qScript cDNA Synthesis Kit (Quantbio, Beverly, USA).

### Metabolite extraction

Soluble metabolites were extracted from freeze dried material. Pulverized material was extracted twice 80% EtOH at 80°C for 1h while shaking. Metabolite extracts were combined and vaporized in a vacufuge concentrator (Eppendorf, Hamburg, Germany). Evaporated pellets were resolved in _dd_H_2_O. Pellets remaining from extraction were washed with 80% EtOH and _dd_H_2_O for starch isolation. 200μl _dd_H_2_O were added to the pellet and samples were autoclaved for 40 min at 121°C. For hydrolytic cleavage 200μl enzyme mix (5 U α-Amylase; 5 U Amyloglucosidase; 200 mM Sodium-Acetate; pH 4.8) were added to the pellet and incubated at 37°C for four hours. Enzymatic digestion was terminated by heating to 95°C for 10 minutes. After centrifugation the supernatant was used for starch quantification.

### Sugar and starch quantification

Sugars (glucose, fructose, sucrose) and hydrolyzed starch concentrations were measured using a NAD^+^-coupled enzymatic assay as described by (Stitt et al., 1989).

### Ion chromatography measurements

Anions, cations, organic acids, sugar alcohols, galactinol and raffinose were measured in a 761 Compact IC system (Metrohm, Herisau, Switzerland). For anion concentration measurements a Metrosep A Supp 4-250/4.0 column and a Metrosep A Supp 4/5 Guard/4.0 guard column (both Metrohm, Herisau, Switzerland) was used. 50mM H_2_SO_4_ was used as anti-ion and 1,8 mM Na_2_CO_3_ together with 1,7 mM NaHCO_3_ dissolved in ultrapure water were used as eluent for anion measurement. For determination of cation concentrations, a Metrosep C4 150/4.0 column and a Metrosep C4 Guard/4.0 guard column (both Metrohm, Herisau, Switzerland) were used. The eluent consisted of 2 mM HNO_3_ and 1,6 mM dipicolinic acid dissolved in ultrapure water. Organic acids were separated via the Metrosep organic acids 250/7.8 column, a Metrosep Organic Acids Guard/4.6 guard column (both Metrohm, Herisau, Switzerland), with 0,25 mM H_2_SO_4_ dissolved in ultrapure water used as eluent and 10mM LiCl as anti-ion. Sugar alcohol, raffinose and galactinol quantification was done by ion chromatography on a Metrosep Carb 2 −250/4.0 column using an 871 IC compact device (Metrohm-Switzerland, Herisau, Switzerland) followed by amperometric quantification. NaOH (0.1 M) with Sodium acetate (10mM) was used as the mobile phase. For peak analysis the corresponding software Metrodata IC Net 2.3 SR5 by Metrohm (Herisau, Switzerland) was used.

### High-performance liquid chromatography measurements

Amino acid concentrations were measured via high performance liquid chromatography in a Dionex (Dionex Softron, Germering, Germany) system, consisting of a Dionex ASI-100 automated sample injector, a Dionex P680 HPLC pump and a Dionex RF2000 fluorescence detector. An AminoPac^®^ PA1 column (Dionex Softron, Germering, Germany) was used for separation of amino acids. 0.1 M NaAc, 7 mM Triethanolamine pH 5.2 was used as eluent. Samples were prepared for measurement by adding 60 μl boric acid buffer (0.2 M; pH 8.8) and 20 μl 6-aminoquinolyl-N-hydroxysuccinimidyl carbamate (3 mg dissolved in 1.5ml Acetonitrile). Samples were vortexed and incubated at 55°C for 10 minutes. Peaks were analyzed using the Chromeleon software (Thermo Fisher Scientific, Waltham, Massachusetts, USA).

### Histological staining, microscopy and determination of cell size

Longitudinal- and cross-sections of sugar beet taproot tissue with a thickness of about 0.5 mm were made with the help of a truffle slicer. Sections were analyzed using a Leica MZ 10 F Binocular (Leica Microsystems, Wetzlar, Germany). For histological staining of cell walls, cross sections were stained using a Fuchsine-Chryosidine-Astra Blue (FCA) staining solution (Morphisto, Frankfurt am Main, Germany). Sections were incubated in staining solution for 5 minutes, washed with water and staining was differentiated in 70% ethanol.

For tracking of Phloem unloading in the root tissue, CFDA (5(6)-Carboxyfluorescein diacetate) was loaded onto source leaves as described by Mehdi et al., 2019. Briefly, the cuticle of the upper surface of source leaves was gently scratched with smooth sand paper. CFDA was prepared freshly from stock (stock: 10 mg ml^−1^ in acetone), by 1:7 dilution (v/v) with ddH_2_O and was pipetted on the roughened leaf surface, which was covered with transparent foil to prevent evaporation through the scratched cuticula. CFDA is non-fluorescent, but membrane permeable before the ester bonds are cleaved to yield the green fluorescent CF (carboxyfluorescein), which then is trapped within cells. Approximately 24h after loading to the leaves, cross sections of taproot / pith tissue were cut using a vibratome (125 μm thickness), or by hand, cell walls were (counter-) stained with propidium iodide and signals detected upon excitation with 488 nm on a Leica SP5 confocal microscope. (Leica, Weimar, Germany).

Mean sizes of pith- and storage parenchyma cells were measured with ImageJ (Rasband, 1997-2018) using microscope pictures of sugar beet cross-sections. For determination of pith cell size, cells from the middle of the pith were chosen. For each cell type, ten cells of typical appearance were measured per section. Overall, seven cross-sections were analyzed for cell size determination.

### ROS staining and Antioxidant measurements

H_2_O_2_ and O_2^-^_ were detected in longitudinal taproot sections according to (Fryer et al., 2002). Sections were vacuum infiltrated with 5mM DAB-HCl, pH3, or 5mM NBT in 50mM potassium-Phosphate-buffer, pH7.8, for 1 hour and incubated at 4°C overnight until pictures were taken the next day. Total and reduced ascorbate concentrations were measured according to Gillespie and Ainsworth (2007). Freshly harvested and pulverized material was dissolved in 6% TCA and immediately used for colorimetric detection of ascorbate at 525nm after the reaction with α-α-bipyridyl. Oxidized DHA concentration was calculated by subtraction of the reduced ascorbate from the total ascorbate concentration. Total and oxidized glutathione concentrations were measured by the glutathione reductase mediated reduction of 5’5-Dithiobis (-2-Nitrobenzoic acid) (DTNB) at 412nm according to (Queval and Noctor, 2007). Reduced glutathione concentration was calculated by subtraction of the oxidized glutathione from the total glutathione concentration.

### Phylogeny

Phylogeny of GOLS and RS isoforms was calculated using the www.phylogeny.fr “one-click” mode (Dereeper et al., 2008). For graphical representation, phylogeny analysis from the Phylogeny.fr platform was loaded into FigTree v1.4.4.

### Figure preparation

Violin plots were generated using BoxPlotR web page (Spitzer et al., 2014). Figures were prepared using Microsoft Excel and Microsoft PowerPoint.

### Statistical analyses

Statistical analysis was performed using Origin Version 2018b (OriginLab Corporation, Northampton, MA, USA). PC analysis was performed using MetaboAnalyst 4.0 web server https://www.metaboanalyst.ca/ (Chong and Xia, 2018).

## SUPPLEMENTAL INFORMATION

**Supplemental Figure S1.** NBT staining of sugar beet taproot sections and antioxidant level of storage parenchyma for ROS detection.

**Supplemental Table S1.** Metabolite concentrations of sugar beet leaf, pith and storage parenchyma tissue.

**Supplemental Figure S2.** Differences in metabolite profile changes between sugar beet tissues upon a shift in growth temperature from 20°C to 0°C.

**Supplemental Figure S3.** Concentrations of starch, glucose, fructose and sucrose in leaf, pith and root tissue under control and freezing temperatures.

**Supplemental Figure S4.** Phylogeny of *Beta vulgaris* GOLS and RS isoforms.

**Supplemental Table S2.** Oligonucleotides used in this study.

## ACKNOWLEDGMENTS

The authors would like to thank Sebastian Stein, Tim Seibel (University of Kaiserslautern), Wiebke Rettberg, Wenke Rettberg and Joern Koch (KWS) for help and support withle sugar beet harvesting and Regina Rhode, Maike Siegel, Ralf Pennther-Hager (University of Kaiserslautern) and Michaela Brock (FAU Erlangen) for excellent technical assistance and plant growth.

