## Supplemental Information for "Cold-triggered induction of ROS- and raffinose-related metabolism in freezing-sensitive taproot tissue of sugar beet"

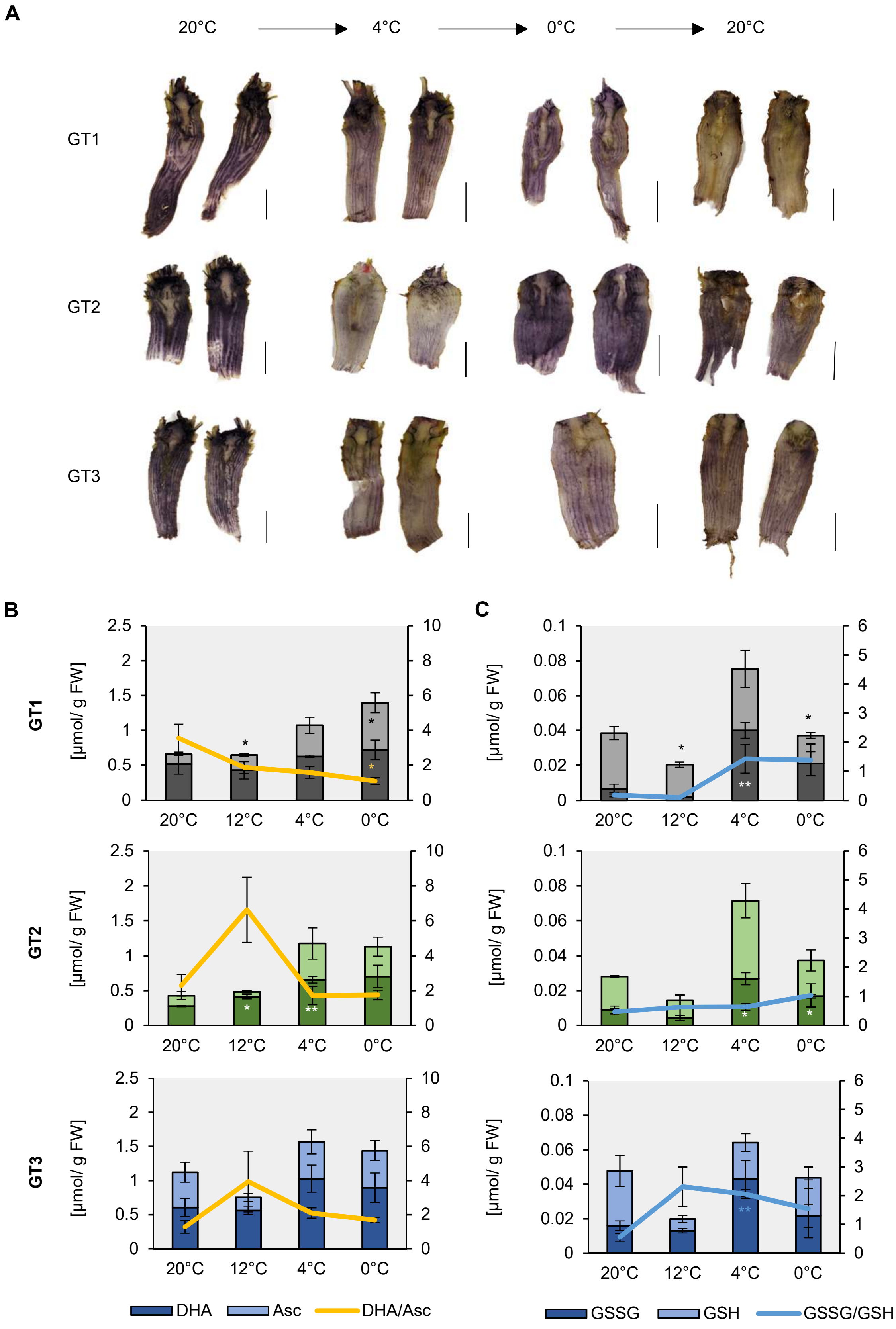


**Supplemental Figure S1: NBT staining of sugar beet taproot sections and antioxidant level of storage parenchyma for ROS detection.** (A) NBT staining is indicative for superoxide accumulation. 10-week old taproots were sectioned and stained after growth under control conditions or at 4°C, 0°C and recovery after freezing at 20°C. Scale bars represent 1cm. (B) and (C) Concentrations of ascorbate (Asc) and dehydro-ascorbate (DHA) (B) or reduced (GSH) and oxidized glutathione (GSSG) (C) in the storage parenchyma of different sugar beet genotypes. Reduced form of the antioxidant depicted in light, oxidized form in dark color. Error bars represent the standard error over the corresponding mean. Asterisks indicate significant differences in concentrations of reduced, or of oxidized antioxidants or of the oxidized/ reduced ratio calculated in relation to the corresponding values at 20°C using Student´s *t*-test (* p< 0,05; ** p< 0.01).

**Supplemental Table S1: Metabolite concentrations of sugar beet leaf, pith and storage parenchyma (root) tissues.** Mean and standard error of measured metabolites in every tissue and temperature analyzed, given in µmol/g DW. Means were calculated of at least three biological replications.

| | | | GT1 | | | | | | GT2 | | | | | | GT3 | | | | | |
|---|---|---|---|---|---|---|---|---|---|---|---|---|---|---|---|---|---|---|---|---|
| | | | leaf | | pith | | root | | leaf | | pith | | root | | leaf | | pith | | root | |
| | | | 20°C | 0°C | 20°C | 0°C | 20°C | 0°C | 20°C | 0°C | 20°C | 0°C | 20°C | 0°C | 20°C | 0°C | 20°C | 0°C | 20°C | 0°C |
| cations | $K^+$ | MEAN | 404.97 | 199.65 | 230.77 | 200.46 | 133.86 | 115.29 | 429.85 | 360.17 | 64.66 | 48.96 | 84.46 | 47.23 | 357.57 | 153.76 | 175.48 | 121.77 | 170.41 | 85.46 |
| | | ±SE | 92.60 | 38.40 | 16.19 | 50.99 | 23.78 | 31.56 | 48.99 | 17.27 | 11.03 | 5.62 | 18.75 | 9.51 | 17.36 | 30.93 | 21.13 | 25.60 | 9.05 | 7.77 |
| | $Na^+$ | MEAN | 622.71 | 414.61 | 123.03 | 130.27 | 41.40 | 55.83 | 686.05 | 284.68 | 30.93 | 18.13 | 15.78 | 7.35 | 854.61 | 515.46 | 102.25 | 43.55 | 53.19 | 17.05 |
| | | ±SE | 118.04 | 63.09 | 20.79 | 28.35 | 10.22 | 18.11 | 105.19 | 5.87 | 8.35 | 4.28 | 3.55 | 3.61 | 88.42 | 51.11 | 20.96 | 6.12 | 16.15 | 2.60 |
| | $NH_4^+$ | MEAN | 28.53 | 25.74 | 17.57 | 33.22 | 7.89 | 10.24 | 37.03 | 23.90 | 6.38 | 7.78 | 2.43 | 3.90 | 40.47 | 24.52 | 9.09 | 17.00 | 7.21 | 8.46 |
| | | ±SE | 2.68 | 4.48 | 3.53 | 5.78 | 2.60 | 1.90 | 4.46 | 2.93 | 1.94 | 0.99 | 0.94 | 0.87 | 4.78 | 1.88 | 1.22 | 5.20 | 1.39 | 0.98 |
| | $Ca_2^+$ | MEAN | 2.74 | 4.32 | 10.82 | 1.96 | 60.78 | 1.13 | 2.80 | 1.67 | 2.64 | 2.27 | 1.14 | 1.58 | 10.06 | 1.03 | 1.51 | 2.08 | 2.06 | 1.07 |
| | | ±SE | 1.07 | 2.96 | 3.90 | 0.63 | 30.76 | 0.16 | 0.81 | 0.83 | 0.45 | 0.32 | 0.27 | 0.60 | 1.66 | 0.29 | 0.27 | 0.36 | 1.36 | 0.57 |
| | $Mg_2^+$ | MEAN | 6.07 | 5.31 | 5.20 | 4.32 | 3.64 | 1.52 | 6.28 | 4.16 | 1.73 | 1.87 | 1.04 | 1.15 | 6.26 | 4.29 | 5.47 | 2.68 | 2.39 | 2.06 |
| | | ±SE | 1.08 | 2.19 | 1.74 | 0.75 | 1.43 | 0.38 | 0.63 | 0.84 | 0.59 | 0.46 | 0.28 | 0.11 | 0.44 | 1.31 | 1.04 | 0.82 | 0.42 | 0.50 |
| anions | $Cl^-$ | MEAN | 420.43 | 935.64 | 163.02 | 180.40 | 51.33 | 58.24 | 487.71 | 302.69 | 35.09 | 27.99 | 18.67 | 14.21 | 309.79 | 552.83 | 123.75 | 78.36 | 89.69 | 30.29 |
| | | ±SE | 57.33 | 229.40 | 29.88 | 13.28 | 8.52 | 14.54 | 207.38 | 25.06 | 13.22 | 3.32 | 6.27 | 6.28 | 68.84 | 108.54 | 33.85 | 15.14 | 24.52 | 6.86 |
| | $NO_3^-$ | MEAN | 213.03 | 197.28 | 342.89 | 234.50 | 58.33 | 113.37 | 271.10 | 46.74 | 21.91 | 8.15 | 16.87 | 6.58 | 150.36 | 85.85 | 118.45 | 71.43 | 56.67 | 20.44 |
| | | ±SE | 91.79 | 52.32 | 75.79 | 49.72 | 16.34 | 37.94 | 82.00 | 6.08 | 6.74 | 1.20 | 8.40 | 3.78 | 19.76 | 26.84 | 23.09 | 16.20 | 10.59 | 1.65 |
| | $PO_4^{3-}$ | MEAN | 10.07 | 15.93 | 24.96 | 15.39 | 25.94 | 26.79 | 18.23 | 25.48 | 2.92 | 6.92 | 11.90 | 21.73 | 9.37 | 18.72 | 0.56 | 10.02 | 5.67 | 26.70 |
| | | ±SE | 3.87 | 4.68 | 10.42 | 7.87 | 3.97 | 5.13 | 4.69 | 4.54 | 0.30 | 1.64 | 2.81 | 0.82 | 0.71 | 7.96 | 0.13 | 2.65 | 2.85 | 4.17 |
| | $SO_4^{2-}$ | MEAN | 13.66 | 34.32 | 7.97 | 8.01 | 5.07 | 3.53 | 42.07 | 11.13 | 1.65 | 3.63 | 1.42 | 3.85 | 15.75 | 24.77 | 1.62 | 5.54 | 2.16 | 4.74 |
| | | ±SE | 2.78 | 6.14 | 0.89 | 2.27 | 0.57 | 0.38 | 19.39 | 2.19 | 0.31 | 0.69 | 0.18 | 0.64 | 3.11 | 5.46 | 0.24 | 1.56 | 0.71 | 0.88 |
| sugars | glucose | MEAN | 35.08 | 137.65 | 9.32 | 14.94 | 3.72 | 1.75 | 60.67 | 256.87 | 8.18 | 24.61 | 2.05 | 2.93 | 46.11 | 113.36 | 9.07 | 9.35 | 3.34 | 3.22 |
| | | ±SE | 8.04 | 12.86 | 2.12 | 5.29 | 0.83 | 0.42 | 15.18 | 40.76 | 1.28 | 8.47 | 0.46 | 0.50 | 7.73 | 26.90 | 1.52 | 0.85 | 0.94 | 0.83 |
| | fructose | MEAN | 36.32 | 216.76 | 2.63 | 12.00 | 3.51 | 1.39 | 46.56 | 348.83 | 4.25 | 8.77 | 2.03 | 1.38 | 23.06 | 142.90 | 4.22 | 3.67 | 3.58 | 2.36 |
| | | ±SE | 5.99 | 24.42 | 0.50 | 6.04 | 0.99 | 0.18 | 20.61 | 48.24 | 0.88 | 2.71 | 0.41 | 0.25 | 6.93 | 33.58 | 0.62 | 0.45 | 0.66 | 0.78 |
| | sucrose | MEAN | 223.75 | 359.05 | 2934.43 | 2428.60 | 3425.73 | 2420.90 | 366.87 | 239.28 | 2335.22 | 2882.36 | 3416.19 | 3544.75 | 383.55 | 309.53 | 2528.06 | 1610.51 | 2850.65 | 3308.58 |
| | | ±SE | 55.18 | 42.83 | 206.75 | 153.34 | 277.17 | 769.00 | 41.16 | 36.77 | 64.86 | 88.03 | 157.12 | 402.09 | 59.30 | 29.16 | 69.42 | 626.46 | 187.24 | 242.89 |
| | raffinose | MEAN | 0.08 | 7.05 | 0.77 | 0.84 | 0.87 | 1.25 | 0.13 | 5.14 | 1.51 | 2.16 | 0.97 | 1.35 | 0.31 | 3.38 | 0.90 | 1.75 | 1.11 | 1.09 |
| | | ±SE | 0.03 | 0.84 | 0.10 | 0.07 | 0.14 | 0.14 | 0.08 | 0.72 | 0.32 | 0.44 | 0.12 | 0.11 | 0.02 | 0.20 | 0.17 | 0.25 | 0.16 | 0.06 |
| sugar alcohol | inositol | MEAN | 0.39 | 1.84 | 0.09 | 0.16 | 0.16 | 0.13 | 0.84 | 1.76 | 0.35 | 0.23 | 0.15 | 0.13 | 0.91 | 1.41 | 0.17 | 0.20 | 0.24 | 0.14 |
| | | ±SE | 0.05 | 0.39 | 0.02 | 0.07 | 0.03 | 0.02 | 0.10 | 0.27 | 0.04 | 0.04 | 0.04 | 0.03 | 0.19 | 0.10 | 0.01 | 0.02 | 0.04 | 0.02 |
| | sorbitol | MEAN | 0.27 | 0.22 | 0.15 | 0.18 | 0.18 | 0.07 | 0.49 | 0.20 | 0.16 | 0.07 | 0.18 | 0.10 | 0.40 | 0.38 | 0.12 | 0.05 | 0.17 | 0.09 |
| | | ±SE | 0.08 | 0.06 | 0.02 | 0.01 | 0.02 | 0.02 | 0.19 | 0.02 | 0.03 | 0.01 | 0.03 | 0.02 | 0.04 | 0.02 | 0.01 | 0.01 | 0.06 | 0.01 |
| | mannitol | MEAN | n.d. | n.d. | 0.04 | 0.08 | 0.08 | 0.11 | n.d. | n.d. | 0.12 | 0.10 | 0.06 | 0.13 | n.d. | n.d. | 0.11 | 0.11 | 0.11 | 0.12 |
| | | ±SE | n.d. | n.d. | 0.00 | 0.05 | 0.01 | 0.02 | n.d. | n.d. | 0.02 | 0.01 | 0.01 | 0.01 | n.d. | n.d. | 0.02 | 0.01 | 0.01 | 0.01 |
| | galactinol | MEAN | 4.90 | 11.65 | 2.91 | 6.01 | 1.83 | 2.44 | 8.17 | 20.70 | 3.20 | 4.39 | 1.46 | 2.30 | 5.80 | 11.13 | 2.63 | 4.70 | 1.58 | 2.83 |
| | | ±SE | 0.81 | 1.56 | 0.38 | 0.68 | 0.16 | 0.31 | 0.55 | 3.18 | 0.26 | 0.43 | 0.18 | 0.19 | 1.28 | 0.38 | 0.19 | 0.49 | 0.39 | 0.19 |
| | starch | MEAN | 100.86 | 9.16 | 8.69 | 9.56 | 12.34 | 16.34 | 73.25 | 6.70 | 7.87 | 4.37 | 7.82 | 10.12 | 166.15 | 3.94 | 9.25 | 3.40 | 8.83 | 15.73 |
| | | ±SE | 28.91 | 2.01 | 2.81 | 1.07 | 3.24 | 1.10 | 16.52 | 0.48 | 1.46 | 0.59 | 3.01 | 1.73 | 36.04 | 0.48 | 2.28 | 0.55 | 4.66 | 2.24 |
| organic acids | citrate | MEAN | 16.44 | 13.59 | 5.17 | 5.92 | 9.03 | 6.94 | 20.84 | 19.15 | 2.01 | 1.74 | 4.56 | 3.10 | 13.84 | 10.89 | 3.37 | 3.50 | 9.91 | 5.54 |
| | | ±SE | 2.94 | 2.73 | 1.13 | 1.16 | 2.08 | 0.80 | 4.67 | 2.31 | 0.32 | 0.35 | 1.68 | 0.71 | 1.01 | 1.96 | 0.37 | 0.52 | 1.68 | 1.35 |
| | malate | MEAN | 22.29 | 6.72 | 1.21 | 2.79 | 2.30 | 2.07 | 24.27 | 15.54 | 1.07 | 1.93 | 1.30 | 1.19 | 17.31 | 10.58 | 0.78 | 1.83 | 2.78 | 1.98 |
| | | ±SE | 2.71 | 1.68 | 0.13 | 0.45 | 0.70 | 0.09 | 7.43 | 3.01 | 0.12 | 0.22 | 0.55 | 0.07 | 3.25 | 1.30 | 0.06 | 0.27 | 0.55 | 0.67 |
| | succinate | MEAN | 1.51 | 0.71 | 0.43 | 0.27 | 0.37 | 0.21 | 0.70 | 0.61 | 0.27 | 0.12 | 0.49 | 0.08 | 0.80 | 0.57 | 0.30 | 0.22 | 0.29 | 0.15 |
| | | ±SE | 0.34 | 0.24 | 0.08 | 0.08 | 0.07 | 0.05 | 0.27 | 0.20 | 0.05 | 0.03 | 0.29 | 0.01 | 0.05 | 0.06 | 0.03 | 0.02 | 0.06 | 0.05 |
| amino acids | aspartate | MEAN | 2.08 | 1.66 | 2.11 | 0.80 | 2.72 | 1.44 | 1.95 | 1.83 | 1.94 | 1.18 | 2.33 | 1.27 | 3.74 | 3.18 | 1.58 | 0.67 | 2.57 | 1.63 |
| | | ±SE | 0.55 | 0.58 | 0.39 | 0.11 | 0.32 | 0.14 | 0.36 | 0.07 | 0.13 | 0.08 | 0.15 | 0.01 | 1.21 | 0.29 | 0.12 | 0.10 | 0.31 | 0.21 |
| | glutamate | MEAN | 6.45 | 2.04 | 3.48 | 2.11 | 2.48 | 1.80 | 5.24 | 1.87 | 2.57 | 3.24 | 1.63 | 2.14 | 8.95 | 2.82 | 2.62 | 2.20 | 2.21 | 2.10 |
| | | ±SE | 0.73 | 0.76 | 0.19 | 0.26 | 0.11 | 0.14 | 0.56 | 0.28 | 0.18 | 0.40 | 0.14 | 0.11 | 1.76 | 0.23 | 0.19 | 0.29 | 0.35 | 0.34 |
| | serine | MEAN | 1.65 | 2.03 | 0.89 | 3.21 | 0.71 | 1.07 | 1.34 | 3.51 | 0.61 | 2.50 | 0.45 | 1.22 | 2.47 | 2.79 | 0.56 | 2.91 | 0.92 | 1.84 |
| | | ±SE | 0.44 | 0.34 | 0.13 | 0.71 | 0.13 | 0.17 | 0.27 | 0.25 | 0.13 | 0.30 | 0.02 | 0.06 | 0.35 | 0.25 | 0.05 | 0.80 | 0.16 | 0.13 |
| | glycine | MEAN | 0.22 | 0.14 | 0.35 | 1.10 | 0.27 | 0.29 | 0.13 | 0.73 | 0.18 | 0.35 | 0.06 | 0.18 | 0.19 | 0.35 | 0.17 | 0.48 | 0.16 | 0.24 |
| | | ±SE | 0.06 | 0.04 | 0.07 | 0.19 | 0.07 | 0.05 | 0.02 | 0.12 | 0.03 | 0.03 | 0.00 | 0.03 | 0.06 | 0.02 | 0.03 | 0.15 | 0.02 | 0.03 |
| | glutamine | MEAN | 0.93 | 0.65 | 2.47 | 4.30 | 1.42 | 1.15 | 0.54 | 0.59 | 1.45 | 2.65 | 0.35 | 1.23 | 1.04 | 1.32 | 1.03 | 3.62 | 0.93 | 2.06 |
| | | ±SE | 0.15 | 0.23 | 0.36 | 0.61 | 0.29 | 0.20 | 0.03 | 0.05 | 0.35 | 0.47 | 0.02 | 0.10 | 0.33 | 0.25 | 0.08 | 1.03 | 0.18 | 0.29 |
| | cysteine | MEAN | n.d. | n.d. | n.d. | n.d. | n.d. | n.d. | n.d. | n.d. | n.d. | n.d. | n.d. | n.d. | n.d. | n.d. | n.d. | n.d. | n.d. | n.d. |
| | | ±SE | n.d. | n.d. | n.d. | n.d. | n.d. | n.d. | n.d. | n.d. | n.d. | n.d. | n.d. | n.d. | n.d. | n.d. | n.d. | n.d. | n.d. | n.d. |
| | histidine | MEAN | 0.17 | 0.23 | 0.11 | 0.28 | 0.07 | 0.04 | 0.08 | 0.53 | 0.09 | 0.37 | 0.03 | 0.08 | 0.10 | 0.36 | 0.07 | 0.26 | 0.04 | 0.08 |
| | | ±SE | 0.10 | 0.10 | 0.02 | 0.05 | 0.02 | 0.01 | 0.01 | 0.03 | 0.02 | 0.05 | 0.00 | 0.00 | 0.03 | 0.02 | 0.01 | 0.10 | 0.01 | 0.02 |
| | threonine | MEAN | 0.56 | 0.67 | 0.33 | 0.54 | 0.20 | 0.15 | 0.50 | 1.34 | 0.23 | 0.32 | 0.13 | 0.16 | 0.65 | 0.91 | 0.22 | 0.51 | 0.24 | 0.30 |
| | | ±SE | 0.17 | 0.23 | 0.05 | 0.12 | 0.03 | 0.03 | 0.06 | 0.09 | 0.06 | 0.04 | 0.01 | 0.01 | 0.21 | 0.08 | 0.01 | 0.18 | 0.04 | 0.03 |
| | alanine | MEAN | 1.90 | 1.19 | 0.66 | 3.40 | 0.57 | 0.69 | 1.33 | 2.12 | 0.38 | 1.35 | 0.26 | 0.40 | 1.67 | 1.76 | 0.45 | 1.82 | 0.72 | 0.77 |
| | | ±SE | 0.24 | 0.35 | 0.10 | 0.94 | 0.17 | 0.07 | 0.23 | 0.11 | 0.08 | 0.38 | 0.05 | 0.05 | 0.34 | 0.14 | 0.06 | 0.58 | 0.17 | 0.09 |
| | arginine | MEAN | 0.56 | 1.38 | 0.08 | 0.20 | 0.03 | 0.02 | 0.59 | 2.16 | 0.02 | 0.09 | 0.02 | 0.01 | 0.65 | 1.45 | 0.04 | 0.10 | 0.04 | 0.03 |
| | | ±SE | 0.41 | 0.79 | 0.03 | 0.08 | 0.01 | 0.01 | 0.33 | 0.32 | 0.01 | 0.02 | 0.00 | 0.00 | 0.38 | 0.25 | 0.01 | 0.03 | 0.01 | 0.01 |
| | proline | MEAN | 0.47 | 1.09 | 0.48 | 1.10 | 0.36 | 0.33 | 0.30 | 2.03 | 0.14 | 0.47 | 0.10 | 0.35 | 0.44 | 0.86 | 0.29 | 0.73 | 0.15 | 0.35 |
| | | ±SE | 0.02 | 0.27 | 0.09 | 0.19 | 0.09 | 0.09 | 0.03 | 0.30 | 0.03 | 0.04 | 0.01 | 0.07 | 0.14 | 0.08 | 0.03 | 0.25 | 0.02 | 0.04 |
| | tyrosine | MEAN | 0.43 | 0.57 | 1.26 | 1.02 | 0.45 | 0.37 | 0.25 | 1.13 | 0.84 | 0.72 | 0.36 | 0.41 | 0.44 | 0.68 | 0.81 | 1.09 | 0.74 | 0.62 |
| | | ±SE | 0.06 | 0.30 | 0.15 | 0.20 | 0.06 | 0.10 | 0.02 | 0.16 | 0.21 | 0.14 | 0.06 | 0.02 | 0.12 | 0.10 | 0.04 | 0.31 | 0.08 | 0.07 |
| | valine | MEAN | 0.61 | 1.26 | 0.38 | 0.88 | 0.27 | 0.33 | 0.40 | 2.09 | 0.17 | 0.61 | 0.19 | 0.33 | 0.65 | 1.49 | 0.23 | 0.62 | 0.33 | 0.41 |
| | | ±SE | 0.31 | 0.74 | 0.11 | 0.14 | 0.05 | 0.07 | 0.07 | 0.25 | 0.03 | 0.09 | 0.02 | 0.02 | 0.27 | 0.19 | 0.04 | 0.21 | 0.05 | 0.06 |
| | methionine | MEAN | n.d. | n.d. | 0.10 | 0.15 | 0.05 | 0.06 | n.d. | n.d. | 0.03 | 0.11 | 0.03 | 0.03 | n.d. | n.d. | 0.04 | 0.18 | 0.05 | 0.03 |
| | | ±SE | n.d. | n.d. | 0.04 | 0.02 | 0.01 | 0.02 | n.d. | n.d. | 0.00 | 0.02 | 0.00 | 0.00 | n.d. | n.d. | 0.01 | 0.05 | 0.01 | 0.01 |
| | isoleucine | MEAN | 0.41 | 0.82 | 0.84 | 1.57 | 0.43 | 0.42 | 0.28 | 1.39 | 0.45 | 1.17 | 0.37 | 0.54 | 0.45 | 0.78 | 0.50 | 1.28 | 0.51 | 0.63 |
| | | ±SE | 0.19 | 0.39 | 0.19 | 0.24 | 0.08 | 0.08 | 0.03 | 0.17 | 0.10 | 0.15 | 0.04 | 0.02 | 0.16 | 0.13 | 0.07 | 0.35 | 0.05 | 0.06 |
| | leucine | MEAN | 0.61 | 1.06 | 1.07 | 1.40 | 0.37 | 0.45 | 0.45 | 2.22 | 0.54 | 1.01 | 0.33 | 0.58 | 0.59 | 1.23 | 0.65 | 1.34 | 0.53 | 0.72 |
| | | ±SE | 0.32 | 0.65 | 0.25 | 0.28 | 0.06 | 0.12 | 0.06 | 0.22 | 0.13 | 0.12 | 0.04 | 0.00 | 0.19 | 0.25 | 0.07 | 0.38 | 0.07 | 0.07 |
| | lysine | MEAN | 0.05 | 0.17 | 0.23 | 0.06 | 0.15 | 0.06 | 0.08 | 0.47 | 0.10 | 0.06 | 0.15 | 0.07 | 0.13 | 0.39 | 0.19 | 0.06 | 0.17 | 0.06 |
| | | ±SE | 0.01 | 0.08 | 0.04 | 0.03 | 0.02 | 0.02 | 0.01 | 0.05 | 0.01 | 0.02 | 0.02 | 0.01 | 0.04 | 0.04 | 0.01 | 0.02 | 0.02 | 0.01 |
| | phenylalanine | MEAN | 0.27 | 0.63 | 0.15 | 0.23 | 0.05 | 0.08 | 0.21 | 1.14 | 0.09 | 0.27 | 0.06 | 0.12 | 0.41 | 0.87 | 0.11 | 0.20 | 0.07 | 0.14 |
| | | ±SE | 0.14 | 0.43 | 0.06 | 0.06 | 0.01 | 0.02 | 0.02 | 0.09 | 0.02 | 0.05 | 0.01 | 0.01 | 0.20 | 0.12 | 0.03 | 0.06 | 0.01 | 0.04 |

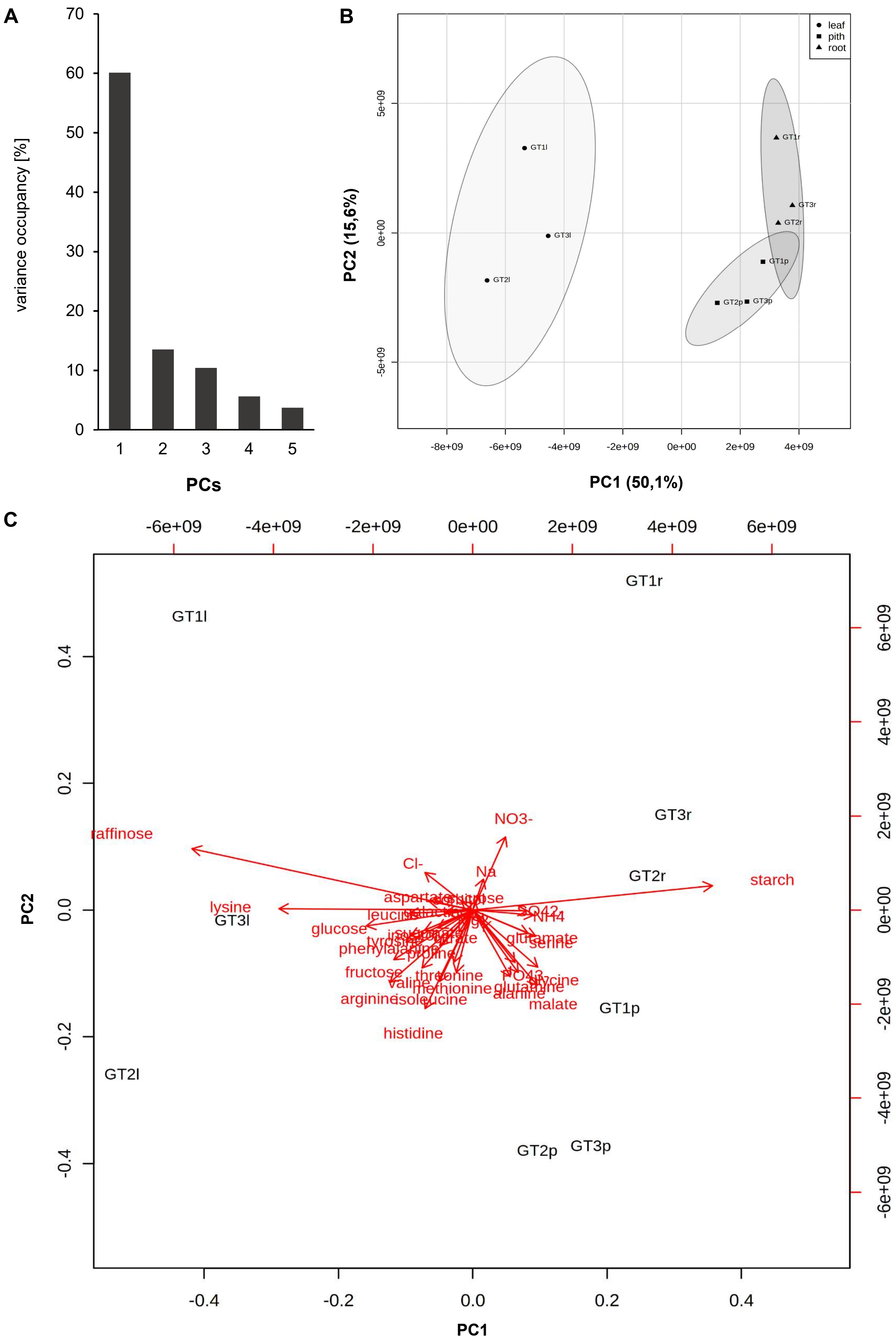


**Supplemental Figure S2: Differences in metabolite profile changes among tissues from three sugar beet cultivars upon a shift in growth temperature from 20°C to 0°C**. Differences were analyzed by Principal Component Analysis. (A) Explained variances of the first five principal components (PCs). (B) Principal component analysis (PCA) of metabolite log2 fold changes after shift in temperature from 20°C to 0°C in different sugar beet tissues. (C) The corresponding loading plot includes the names of 36 metabolites contributing to the separation of PC1 and PC2. The means of metabolite concentrations of three biological replications for each tissue and genotype measured in plants grown at 20°C and 0°C were used to calculate the log2 fold changes 20°C/0°C and used as loadings for this analysis.

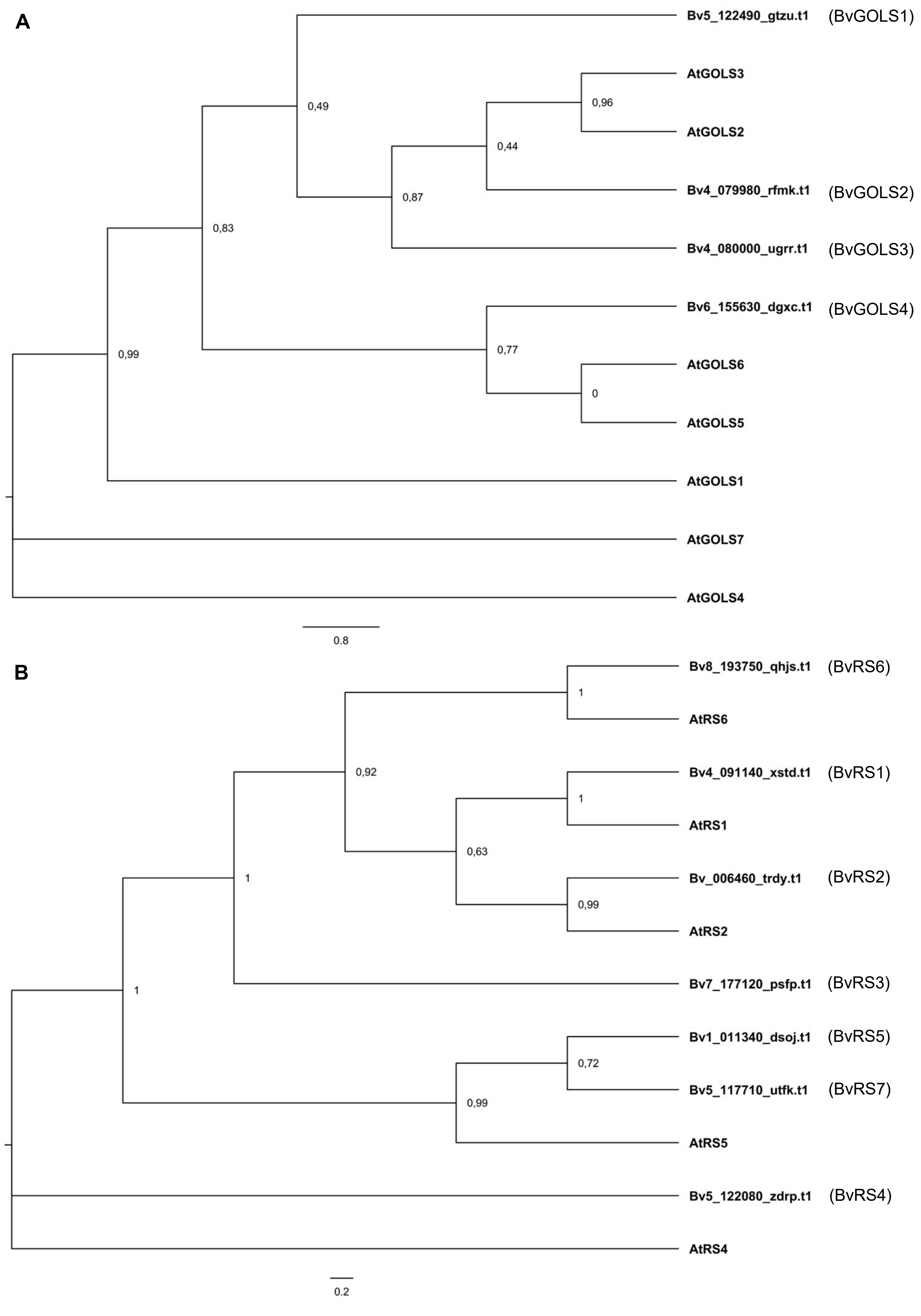


**Supplemental Figure S3: Phylogeny of *Beta vulgaris* GOLS and RS isoforms.** (A) Phylogenetic tree of galactinol synthase amino acid sequences from sugar beet and *Arabidopsis thaliana*. Sugar beet proteins are named according to their identifiers. Arabidopsis proteins had the following identifiers: AtGOLS1: AT2G47180; AtGOLS2: AT1G56600; AtGOLS3: AT1G09350; AtGOLS4: AT1G60470; AtGOLS5: AT5G23790; AtGOLS6: AT4G26250; AtGOLS7: AT1G60450. (B) Phylogenetic tree of raffinose synthase amino acid sequences from sugar beet and *Arabidopsis thaliana*. Sugar beet proteins are named according to their identifiers. Arabidopsis proteins had the following identifiers: AtRS1: AT1G55740; AtRS2: AT3G57520; AtRS4: AT4G01970; AtRS5: AT5G40390; AtRS6: AT5G20250. Phylogenetic trees were calculated using the “one-click” mode of http://www.phylogeny.fr and visualized using FigTree v1.4.4. Branch labels represent branch support values

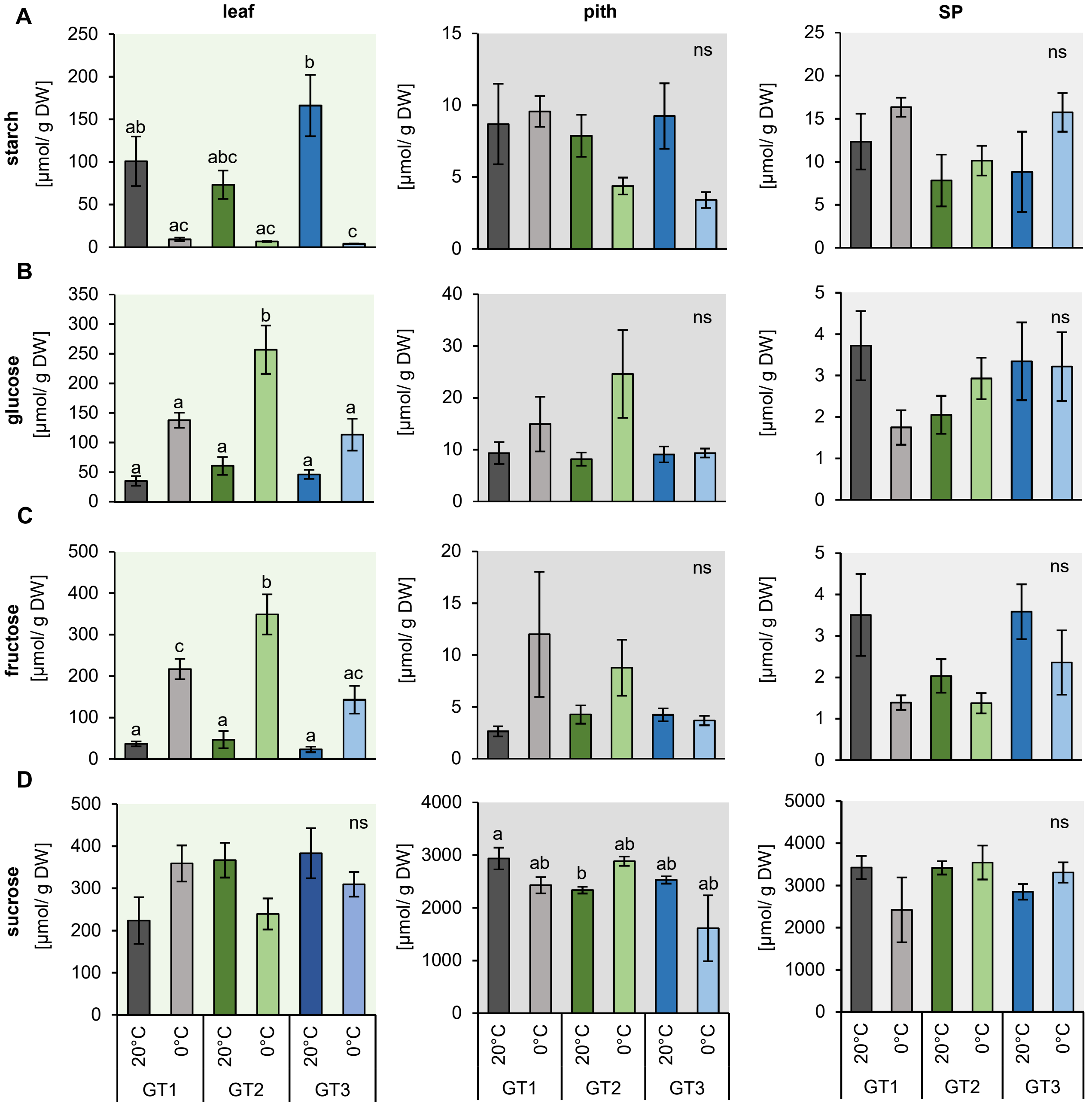


**Supplemental Figure S4: Concentrations of starch, glucose, fructose and sucrose in leaf, pith and storage root tissue under control and freezing temperatures.** Metabolites were measured in plants grown at 20°C and plants transferred to 12°C, 4°C and harvested at 0°C soil temperature. Plants were dissected in the three different tissues leaf, pith and root. Starch (A), glucose (B) fructose (C) and sucrose (D) values represent the mean of three biological replicates for each of the tested cultivars. Error bars represent the standard error of the corresponding mean. Letters indicate the same level of significance for each measured concentration, calculated via two-way ANOVA corrected with post hoc Bonferroni test with $p < 0.05$.

**Supplemental Table S2: Primer used for the analysis of gene expression in *B. vulgaris* tissues.**
Primer Sequences, melting temperature (Tm), as well as amplicon size and correlation coefficient ($R^2$) used for efficiency testing (E) given for each primer used in the analysis. *BvUGD1* served as a reference gene for transcript normalization.

| Locus | Gene symbol | Primer sequence (5′-3′) | Tm (°C) | $R^2$ | E (%) | Amplicon size (bp) |
|---|---|---|---|---|---|---|
| *BVRB_7g172340* | *UGD1* | F: GTAGTGAAGCAATGCCGAGG | 61 | 0.99 | 105.7 | 120 |
| | | R: AGTCAAGTCTGCTGCTTTGC | 58 | | | |
| *BVRB_4g075130* | *CLV2* | F: ATTGAGGTCTCTGCCGGAAA | 58 | 0.99 | 88.4 | 166 |
| | | R: ATCGTGGCCAACAGAAGAGA | 58 | | | |
| *BVRB_3g066590* | *CBF3* | F: GCACATGATGTAGCTGCGAT | 59 | 1.00 | 76.5 | 184 |
| | | R: TCTACCGCCGTTTCCTCTTT | 59 | | | |
| *BVRB_5g122490* | *GOLS1* | F: TTGGTGAAGAAGTGGTGGGA | 58 | 1.00 | 96.1 | 159 |
| | | R: AGCAGCAGAAGGAGCAGTAA | 55 | | | |
| *BVRB_4g079980* | *GOLS2* | F: CTGAGGACAAGTTAGGCCCA | 64 | 0.99 | 88.8 | 229 |
| | | R: AATGTTTTCAGGGTGACGCC | 66 | | | |
| *BVRB_4g080000* | *GOLS3* | F: CGCATTTGGGAGTTTGTGGA | 68 | 1.00 | 118.0 | 201 |
| | | R: GCCAGCACACTCTATTTGGG | 64 | | | |
| *BVRB_006460* | RS2 | F: GCAGCATCTCATTTACGCCA | 58 | 1.00 | 104.3 | 203 |
| | | R: AAACCGGCCACATCCTCTTA | 58 | | | |
| *BVRB_1g011340* | *RS5* | F: AGCCATTCCCCATCAAAGGA | 67 | 0.99 | 105.9 | 170 |
| | | R: CAAATGCGATCGACTCCCAG | 67 | | | |
| *BVRB_8g189640* | *ZAT10* | F: AACATACAAGTGCGGCGTTT | 56 | 0.99 | 90.2 | 186 |
| | | R: GCAGATTGAGCAGACGTGAG | 61 | | | |
| *BVRB_2g037310* | *ZAT12* | F: GCCAATTTGCCTCCTTCCAA | 58 | 0.99 | 99.8 | 180 |
| | | R: TATGACCTCCCAACGCTTGT | 58 | | | |
| *BVRB_9g206190* | *SOD* | F: ATTCTCGCTTCCCACCCTAC | 61 | 0.99 | 89.2 | 179 |
| | | R: AATGGTGAGGGGTTTAGGGG | 61 | | | |
| *BVRB_1g021080* | *CAT* | F: GGCTGGCAAAGTACACTACG | 61 | 1.00 | 82.1 | 242 |
| | | R: TCCTCAGGCCATGTCTTTGT | 58 | | | |
| *BVRB_9g207350* | *APX* | F: CGAGAAGGCCAAGAGAAAGC | 61 | 0.98 | 74.7 | 200 |
| | | R: GGCTCCAACAACCTAACAGC | 61 | | | |
| *BVRB_7g159920* | *MDAR* | F: AAAACTGTCGTGGTTGGTGG | 65 | 0.99 | 91.8 | 229 |
| | | R: CCGTCACCTCTCCGTTATCA | 65 | | | |
| *BVRB_2g039390* | *DHAR* | F: CCTTTAAACGTCCGGTGGTC | 61 | 0.99 | 97.9 | 154 |
| | | R: AGCTTGTTCGGAGTGGTGAT | 58 | | | |
| *BVRB_2g044150* | *GPX* | F: CACAGTGTGGGTTGACATCA | 63 | 0.99 | 89.0 | 217 |
| | | R: GGAGCTGTGTTTGAACCGTT | 64 | | | |
| *BVRB_3g069540* | *GR* | F: AGGGCTGTTGTCGCTAGAAA | 58 | 1.00 | 99.7 | 202 |
| | | R: GTTCAACACCAACAGCGTC | 58 | | | |
